# Isoform-Specific Roles of NTRK2 in Pulmonary Vascular Regeneration

**DOI:** 10.1101/2025.05.11.653351

**Authors:** Cheng Tan, Ziyi Liu, Xiangdi Mao, Yunpei Zhang, Xiaolei Li, Nicole Pek, Hailu Fu, Yaping Liu, Vladimir V. Kalinichenko, Gloria S. Pryhuber, Renzhong Lu, Li Lai, Yifei Miao, Minzhe Guo, Mingxia Gu

**Affiliations:** Perinatal Institute, Division of Pulmonary Biology, Cincinnati Children’s Hospital Medical Center, Cincinnati, OH; Center for Stem Cell and Organoid Medicine (CuSTOM), Division of Developmental Biology, Cincinnati Children’s Hospital Medical Center, Cincinnati, OH; Department of Pediatrics, University of Cincinnati School of Medicine, Cincinnati, OH; Department of Anesthesiology and Perioperative Medicine, David Geffen School of Medicine, Eli and Edythe Broad Center for Regenerative Medicine and Stem Cell Biology, University of California, Los Angeles (UCLA), Los Angeles, CA 90095, USA; Department of Biochemistry and Molecular Genetics, Feinberg School of Medicine, Northwestern University, Chicago, IL; Robert H. Lurie Comprehensive Cancer Center of Northwestern University, Chicago, IL; Phoenix Children’s Research Institute, Department of Child Health, University of Arizona College of Medicine-Phoenix, Phoenix, AZ; Division of Neonatology, Phoenix Children’s Hospital, Phoenix, AZ; Division of Neonatology, Department of Pediatrics, University of Rochester Medical Center, Rochester, NY; Department of Cardiovascular Sciences, Houston Methodist Research Institute, Houston, TX, United States

**Keywords:** Bronchopulmonary dysplasia, Vascular injury and repair, Multiome, Alternative splicing, NTRK2 isoform, RBFOX2, iPSC-derived vessel organoids, Lipid nanoparticles, RNA therapy

## Abstract

Bronchopulmonary dysplasia (BPD) is a chronic lung disease in premature infants with no curative therapy, characterized by impaired alveologenesis and capillary formation. However, the molecular mechanisms underlying endothelial dysfunction, a key driver of BPD pathogenesis in human, remain poorly understood. Here, through multiomic profiling of vascular endothelial cells isolated from control and BPD patient lungs, we uncovered an expansion of general capillary endothelial cells (gCap) with aberrant expression of the neurotrophic receptor tyrosine kinase 2 (NTRK2) in BPD. Importantly, we identified a pathological NTRK2 isoform switch that dictates the regenerative capacity of gCap cells. Full-length NTRK2 (NTRK2-FL) promoted gCap regeneration in response to hyperoxic injury, whereas RBFOX2-mediated splicing of NTRK2-FL into a truncated isoform (NTRK2-T1) contributed to maladaptive responses and irreversible alveolar simplification in severe BPD cases. Restoring NTRK2-FL using lipid nanoparticle–delivered mRNA promoted angiogenesis and reversed alveolar simplification in vessel organoids and BPD-like mice. These findings identified NTRK2 isoform imbalance as a key driver of endothelial dysfunction and support isoform-specific RNA therapy as a promising strategy for vascular regeneration and repair.

**HIGHLIGHTS:** • Multiomic and spatial profiling reveal abnormal gCap subtype in human BPD lungs

• NTRK2 isoform switch dictates endothelial regeneration in response to hyperoxic injury

• RBFOX2 mediates splicing from NTRK2-FL to NTRK2-T1, driving irreversible alveolar damage

• NTRK2-FL mRNA therapy restores vascular regeneration after injury

## INTRODUCTION

Bronchopulmonary dysplasia (BPD) is a significant and persistent complication of prematurity, first described in 1967 as a consequence of mechanical ventilation and oxygen toxicity in preterm neonates. Despite advances in neonatal care, BPD remains a leading cause of neonatal morbidity,^1^ particularly among extremely preterm infants born before 28 weeks of gestation.^2,3^ Characterized by impaired alveolarization and abnormal pulmonary vascular development, BPD can lead to lasting respiratory and cardiovascular complications. While interventions such as antenatal corticosteroids and surfactant replacement therapy have mitigated acute respiratory distress, current treatments including oxygen supplementation, non-invasive ventilation, and post corticosteroids, remain largely palliative and fail to address the developmental arrest that defines BPD.^4–6^ For instance, although postnatal dexamethasone can reduce short-term oxygen dependency, it is associated with increased neurodevelopmental risk and fails to improve long-term pulmonary outcomes.^7,8^ Similarly, while oxygen therapy is life-saving, it exacerbates oxidative stress and damages the developing pulmonary vasculature.^9^ As a result, many BPD survivors face lifelong sequelae such as pulmonary hypertension and impaired exercise capacity.^10–12^ These limitations underscore the urgent and unmet need for therapies that target the fundamental mechanisms of BPD and promote alveolar capillary regeneration.

Emerging evidence positions vascular endothelial cell (EC) dysfunction as both a key pathological driver and a potential therapeutic target in BPD. Exposure to hyperoxia and mechanical ventilation in preterm infants disrupts normal pulmonary vascular development, resulting in vascular rarefaction and increased pulmonary vascular resistance, key factors contributing to pulmonary hypertension and exacerbating morbidity and mortality.^13^ The pulmonary vasculature also plays an active role in alveolar maturation via endothelial-epithelial interactions mediated by signaling molecules such as VEGF, PDGF, and nitric oxide.^14–16^ Therefore, its dysfunction can lead to damage of surrounding cell types and impair alveologenesis in infants. Although EC dysfunction such as impaired proliferation and increased endothelial-to-mesenchymal transition (EndoMT) has been associated with BPD pathology,^17,18^ the underlying molecular mechanisms driving these phenotypic changes remain unclear, especially at single cell resolution in human.

Recent single-cell transcriptomic studies revealed a significant reduction of *Ntrk2* expression in alveolar general capillary endothelial cells (gCap) in mouse lungs exposed to hyperoxic injury,^19^ whereas two other studies reported a significant increase of *Ntrk2* in injury-induced capillary endothelial cells (iCAPs) in mice exposed to viral infection or hyperoxic injury,^20^ as well as in activated capillary ECs in response to bleomycin challenge.^21^ This discrepancy is at least partially due to the lack of in-depth analysis of dynamic changes in *Ntrk2* expression across different stages and severities of lung injury, as well as a limited understanding of the distinct NTRK2 isoforms, which we show in current studies, to play fundamentally different roles in regulating pulmonary capillary endothelial cell functions during lung development and in response to injury.

In this study, we investigate the pivotal role of NTRK2 isoform imbalance in pulmonary endothelial dysfunction and impaired regenerative capacity. By integrating single-cell multiomic analyses of human BPD lung tissues with functional assessments in hyperoxia-exposed vessel organoids derived from human iPSCs and murine models with BPD-like phenotypes, we aim to elucidate the distinct roles and dynamic regulation of full-length versus truncated NTRK2 isoforms in governing gCap homeostasis and responses to hyperoxic injury across different stages of disease progression. Understanding this fundamental mechanism will provide a strong rationale for developing endothelial-specific therapies to improve both short- and long-term outcomes in BPD and related conditions characterized by oxygen-induced endothelial injury.

## RESULTS

### Multiomic single cell analysis identified enrichment of NTRK2^+^ gCap-C2 in BPD patients

To investigate the molecular and functional abnormalities of endothelial cells (ECs) in BPD, we performed multiomic single cell analysis on lung samples from 8-21-month-old BPD patients (n=3) and age-matched controls (n=2) **(Table 1)**. Single-cell suspensions were prepared as previously described,^22^ followed by EC (CD31^+^) enrichment, 10x single-nucleus multiome experiment, sequencing, and downstream analysis **(Figure 1A**, **S1A-C)**. We identified 15 distinct cell types, including six endothelial sub-types according to their marker genes: gCap-C1 (*SLC6A4*, *FCN3*), gCap-C2 (*CLIC5*, *NTRK2*), aCap (*HPGD*, *EDNRB*), Arterial (*GJA5*, *DKK2*), Venous (*ACKR1*, *CPE*), and Systemic Venous endothelium (*ABCB1*, *COL15A1*) (**Figure 1B**, **S1D**). Notably, the proportion of gCap-C2 ECs was significantly elevated in BPD samples as compared to controls **(Figure 1C, S1E-F)**. Interestingly, within the gCap population which is known for its proliferative nature as a progenitor cell type in the alveolus, gCap-C1 retained normal expression of proliferation and cell cycle genes, whereas gCap-C2 showed a marked reduction, suggesting functional abnormalities of gCap-C2 in BPD **(Figure S1G-H).** Next, we compared the transcriptomes of gCap- C1 and gCap-C2 and identified *NTRK2* as the most upregulated gene among all differentially expressed genes (DEGs) in gCap-C2 **(Figure 1D-E)**. Notably, *NTRK2* expression was consistently elevated in lung tissues from all three BPD patients compared to controls, as demonstrated by both multiomic single cell analysis **(Figure 1F)** and *in situ* hybridization **(Figure 1G)**.

**Figure 1:**
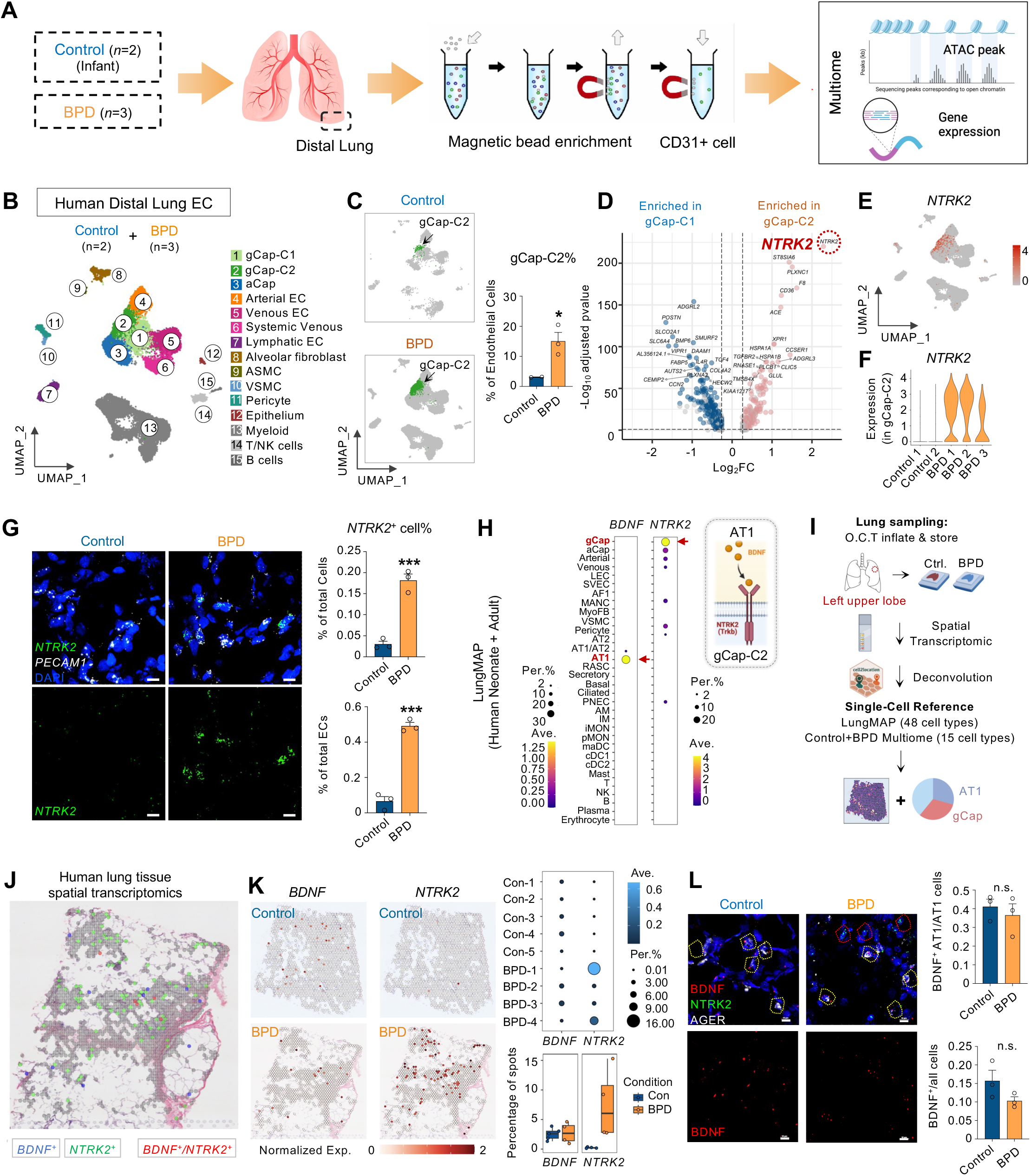
Multiomic single cell analysis revealed the enrichment of general capillary endothelial cells (gCap) C2 subtype and neurotrophic receptor tyrosine kinase 2 (*NTRK2*) in bronchopulmonary dysplasia (BPD) patients (A) Experimental design for comparative 10x single-nucleus multiome RNA+ATAC analyses of CD31 enriched distal lung samples from BPD (ages 8-21 months, n=3) and age-matched control patients (n=2). (B) Uniform manifold approximation and projection (UMAP) visualization of 24,749 cells from (A) after quality control, clustered into 15 cell types, including 6 endothelial subclusters according to specific markers. aCap, aerocyte; ASMC, airway smooth muscle cell; VSMC, vascular smooth muscle cell. (C) UMAP plots showing gCap-C2 cells in control and BPD samples and bar plot demonstrating the higher percentage of gCap-C2 population in BPD lungs compared with controls. Data presented as mean ± SEM, control n=2, BPD n=3. \**P* < 0.05, BPD vs. Control, unpaired two-tailed *t* test. (D) Volcano plot showing gene differential expression between gCap-C2 and gCap- C1 cells. *NTRK2* (red) was enriched in gCap-C2. (E) UMAP presenting RNA expression of *NTRK2* in all samples. (F) Violin plots showing the upregulation of *NTRK2* in gCap-C2 of individual BPD patients compared with controls. (G) Representative images (left) and quantification (right) of *NTRK2^+^* ECs from BPD and control patients using RNA-scope staining. *NTRK2* (green), PECAM-1 (white), and DAPI (blue). Scale bar, 10 μm. Data presented as mean□±□SEM, n=3, \*\*\**P* < 0.001, BPD vs. Control, unpaired two-tailed *t* test . (H) Dot plot presenting the expression of *NTRK2* and its ligand, brain-derived neurotrophic factor (*BDNF*), in each cell type in LungMAP human lung single cell reference (CellRef). (I) Experimental design and analysis of 10x Visium spatial transcriptomics (ST) profiling of BPD and control lung samples. Deconvolution of cell types in the ST data was performed using Cell2location using both the LungMAP CellRef and the present multiomic single cell profiling in (B) as the single cell reference. (J) Demonstration of the spatial patterns of *NTRK2*+, *BDNF*^+^, and *NTRK2*^+^/*BDNF*^+^ spots using ST profiling. (K) ST and dot plot visualization of the expression of *BDNF* and *NTRK2* in BPD and control samples. (L) The expression pattern of BDNF and NTRK2 in BPD and control lung samples using immunofluorescence staining (left), with quantification on the right. NTRK2 (green), BDNF (white), DAPI (blue). Scale bar, 10 μm. Data presented as mean□±□SEM, n=3, unpaired two-tailed *t* test.

**Table 1.** Demographic Information of BPD Patients and Age-matched Controls

**Table 2.** qPCR primers

**Table 3.** mRNA sequence

**Table 4.** Patients information for human lung tissue sections

**Table 5.** Key Resources

As a tyrosine kinase receptor, NTRK2 primarily binds brain-derived neurotrophic factor (BDNF). Interestingly, we found that in human lungs, alveolar type 1 (AT1) cells predominantly secret *BDNF* to target *NTRK2*-expressing gCap ECs, as evidenced by scRNA-seq analysis of neonatal and adult lung tissues **(Figure 1H)**. Spatial transcriptomic profiling of lung samples from both BPD patients and controls **(Figure 1I, S2A-C)** further confirmed the juxtaposed localization of *NTRK2* and *BDNF* in human lung tissues **(Figure 1J)**. Consistent with our multiomic analysis, *NTRK2* expression was significantly elevated in BPD lung samples, whereas the expression of *BDNF,* as well as other two ligands *NTF3* and *NTF4*, remained comparable between BPD and control groups **(Figure 1K, S2D-F)**. This expression pattern of *BDNF* was further validated by RNAScope **(Figure 1L)**.

To further understand the impact of these abnormal gCap-C2 cells on surrounding cell types in the alveolar niche, we systematically identified significant ligand-receptor (L-R) interactions associated with gCap-C2 cells through CellChat analysis on the multiome profiling of the BPD samples (**Figure S3A**). These candidate L-R pairs were subsequently subjected to spatial colocalization analysis using spatial transcriptomics (ST) data. To determine the co-expression probability of ligand- receptor partners, we integrated three-dimensional information: 1) spatial gene expression profiles, 2) spatial neighborhood relationships between tissue spots, and 3) cell type deconvolution results. Specifically, we evaluated whether ligand-receptor pairs co-localized either within individual *NTRK2*^+^ gCap-C2-enriched spots or between adjacent (1-hop) neighboring spots (**Figure S3B**). Comparative analysis between BPD and control lungs revealed L-R pathways originating from gCap-C2 cells, including *SEMA3C*, *EDN1*, *JAM2*, and *LAMA4* signaling networks (**Figure S3C**). Notably, we observed significant upregulation of EDN1 signaling from gCap- C2 cells within BPD specimens (**Figure S3D**).

Collectively, we identified an abnormal gCap state associated with human BPD pathology characterized by elevated *NTRK2* expression, while its ligand BDNF remained unchanged.

### Increase of Truncated NTRK2 Isoform in gCap ECs from Severe BPD Patients

NTRK2 exists in two major isoforms: full-length NTRK2 (NTRK2-FL), which contains an intracellular tyrosine kinase domain, and NTRK2-T1, a truncated isoform lacking tyrosine kinase activity **(Figure 2A).**^23,24^ In the brain, these isoforms have been reported to exhibit distinct functional properties, with NTRK2-T1 playing roles that diverge from those of NTRK2-FL.^25^ During murine lung development, Ntrk2 expression exhibited a distinct temporal pattern, peaking within the first 14 days postnatal specifically in gCap cells, with lower detection across other pulmonary cell populations (**Figure S4A**). Similar, single cell RNA-seq of human lung in different development stages revealed that *NTRK2* expression in gCap cells was restricted exclusively to the neonatal stage (**Figure S4B**).^26^ To further determine the expression patterns of different NTRK2 isoforms in control and BPD lungs in different development stages, we designed RNA-Scope probe specifically recognizing shared domain or tyrosine kinase domain of the *NTRK2* in both mouse and human tissues (**Figure S4C**). Validation of these probes in the human brain cortex confirmed predominant expression of the *NTRK2-FL* isoform (**Figure S4D**), consistent with previous findings.^27,28^ In human lungs, *NTRK2-FL* expression peaked at the saccular and early alveolar stages, then declined to basal levels after 6 months as the lung continued to mature (**Figure S4E-F**). These findings further highlight a critical role for NTRK2-FL in alveologenesis.

**Figure 2:**
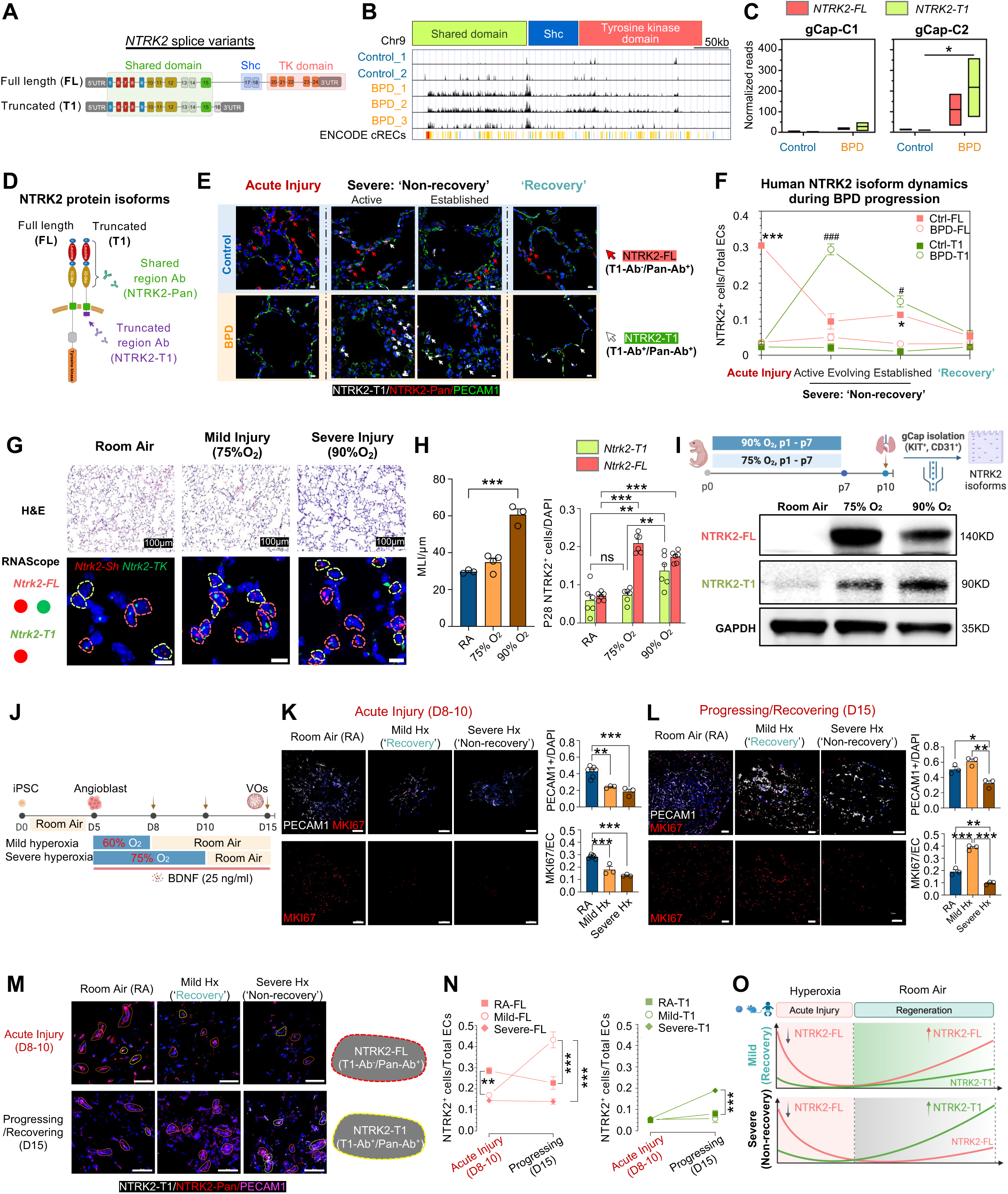
Increased Expression of Truncated *NTRK2* Isoform (*NTRK2-T1*) in Lung gCap ECs from Severe BPD Patients (A) Schematic diagram of structural domains of full length *NTRK2* isoform (*NTRK2- FL*) and *NTRK2-T1*. (B) RNA reads of from the mMultiome profiling presenting mapping *NTRK2* shared domain, Shc region and tyrosine kinase domain infrom gCap cells of each biopsysample. (C) Box plot showing normalized reads of *NTRK2-FL* (red) and *NTRK2-T1*(green) from (B) in gCap-C1 and gCap-C2 in BPD and control groups. Normalized reads of NTRK2-FL were calculated by summing the reads from Shc and tyrosine kinase domain. Normalized reads of NTRK2-T1 were calculated by summing the reads from shared domain minus normalized reads of NTRK2-FL. Data presented as mean (min, max), n=3. *P < 0.05, two-way ANOVA followed by Šídák’s multiple comparisons tests. (D) Demonstration of antibody targeting shared region of NTRK2 (NTRK2-Pan) and antibody targeting truncated region of NTRK2 (NTRK2-T1). (E) The location and expression pattern of NTRK2 isoforms in the distal lungs from different stages of BPD patients, compared to age matched control groups. NTRK2- T1(white), NTRK2-Pan(red), PECAM-1(green), DAPI (blue). Red arrows indicate NTRK2-FL (NTRK2-T1^-^/NTRK2-Pan^+^), and white arrows indicate NTRK2-T1 (NTRK2-T1^-^/NTRK2-Pan^+^). Scale bar, 10 μm. (F) Line plot showing the percentage of different NTRK2 isoform-positive cell among all endothelial cells from (E). Data presented as mean ± SEM, n=3 patients. \**P* < 0.05, \*\*\**P* < 0.001, BPD-FL vs. control-FL, ^#^*P* < 0.05, ^###^*P* < 0.001, BPD-T1 vs. control-T1, one-way ANOVA followed by Bonferroni’s multiple comparisons test. (G) H&E staining (top) and RNA-scope staining (bottom) of lung tissues from postnatal day 28 mice (P28) under different hyperoxia conditions (see STAR method). Scale bar (up), 100 μm. Scale bar (down), 10μm. *Ntrk2-Sh*(red), *Ntrk2-TK* (green). Red circles indicate *Ntrk2-FL* mRNA and green circles indicate *Ntrk2-T1* mRNA. (H) Quantification of MLI (left, n=3) and percentage of *NTRK2*^+^ cells(right, n=6). Data presented by mean ± SEM. \*\**P* < 0.01, \*\*\**P* < 0.001, one-way ANOVA followed by Bonferroni’s multiple comparisons test. (I) Experimental designs for BPD mouse models and gCap isolation(up) and Western blot result of NTRK2-FL, NTRK2-T1 and GAPDH. (J) Experimental design of BPD vessel organoid models under different hyperoxia levels (see STAR method). (K) Representative IF staining images of D8-D10 vessel organoid (left) and quantifications (right). PECAM-1(white), MKI67(red), DAPI (blue). Scale bar, 100μm. Data present by mean ± SEM, n = 3 iPSC lines. \*\**P* < 0.01, \*\*\**P* < 0.001, one-way ANOVA followed by Bonferroni’s multiple comparisons test. (L) Representative IF staining images of D15 vessel organoid (left) and quantifications (right). Scale bar, 100μm. Data present as mean ± SEM, n = 3 iPSC lines. \**P* < 0.05, \*\**P* < 0.01, \*\*\**P* < 0.001, one-way ANOVA followed by Bonferroni’s multiple comparisons test. (M) The locations of NTRK2 isoforms in vessel organoids under hyperoxia treatments. NTRK2-T1(white), NTRK2-Pan(red), and PECAM-1(pink). Red circles indicate NTRK2-FL^+^ cell, yellow circles indicate NTRK2-T1^+^ cell. Scale bar: 100 μm. (N) Line plot showing NTRK2-FL and NTRK2-T1 percentages among total endothelial cell (ECs) from (L). Scale bar, 50 μm. Data presented as mean ± SEM, n = 3 iPSC lines. \*\**P* < 0.01, \*\*\**P* < 0.001, one-way ANOVA followed by Bonferroni’s multiple comparisons test. (O) Demonstration figure showing the dynamic changes of NTRK2 isoforms shared by human sample, mouse model and vessel organoid model.

Surprisingly, the previously observed increase in *NTRK2* in BPD gCap cells was attributed to the truncated isoform *NTRK2-T1*, as evidenced by increased reads mapping to the shared domain but not the tyrosine kinase domain **(Figure 2B-C)**. To further validate this finding at the protein level, we employed two distinct antibodies:^28^ one specifically recognizing the truncated region of NTRK2 (NTRK2-T1 antibody) and another recognizing the shared domain of both NTRK2 isoforms (NTRK2-Pan) **(Figure 2D).** Interestingly, we observed remarkably dynamic changes of NTRK2 isoforms across different stages of BPD progression: at the acute injury stage (≤4 month), NTRK2-FL (T1-Ab^-^/Pan-Ab^+^, red arrow) was significantly downregulated in BPD compared with controls. During the disease progression phase (7-14 month), NTRK2-T1 (T1-Ab^+^/Pan-Ab^+^, white arrow) was significantly increased in BPD cases with severe, non-recovery outcomes, consistent with our multiomic analysis. In recovered BPD cases discharged from the hospital (≥15 month), both NTRK2 isoforms went back to baseline levels **(Figure 2E-F)**. These findings suggest an initial global reduction in NTRK2 expression during acute injury, followed by maladaptive upregulation of NTRK2-T1 during disease progression in severe BPD cases, and normalization of both isoforms to baseline upon recovery.

The etiology of BPD in humans is multifactorial, including mechanical stress, inflammation, and hyperoxia-induced injury. To assess whether *NTRK2* splicing in BPD gCap cells is driven specifically by hyperoxia, and to directly compare mild (75 percent O₂, recovery) and severe (90 percent O₂, non-recovery) cases at matched developmental time points, we used the hyperoxia-induced alveolar simplification model , a well-established mouse model of BPD pathophysiology.^29^ We found that in mild injury group where human tissues were inaccessible, NTRK2-FL was significantly upregulated compared with room air controls **(Figure 2G-H)**, indicating its role in promoting alveolar regeneration and repair. In contrast, the severe group showed increased *Ntrk2-T1* expression, recapitulating the pattern seen in end-stage BPD patients and contributing to impaired regeneration following hyperoxia injury **(Figure 2G-H)**. Western Blot analysis on gCap ECs (KIT^+^/CD31^+^) isolated from the mouse lungs exposed to different levels of hyperoxic injuries confirmed a marked increase in NTRK2-FL in mild cases, and an increase in NTRK2-T1 in severe cases, leading to maladaptive response in the lung **(Figure 2I, S5)**.

To further investigate the role of NTRK2-FL/NTRK2-T1 isoform switching in a human context, we utilized human iPSC-derived vessel organoids exposed to mild and severe hyperoxia conditions **(Figure 2J)**. In the acute injury phase, we observed a significant reduction of proliferative ECs, which directly correlated with the severity of the injury levels **(Figure 2K)**. During the recovery phase, ECs were highly proliferative in the mild injury group, as indicated by increased EC numbers and MKI67 staining, but showed limited proliferation and reduced EC numbers in the severely injured vessel organoids **(Figure 2L)**. These functional changes corresponded with elevated NTRK2-FL expression in vessel organoids exposed to mild hyperoxia, and increased NTRK2-T1 in those subjected to severe injury **(Figure 2M-N)**.

Collectively, through comprehensive analyses of human BPD lung tissues, BPD-like mouse models, and vessel organoids, we found: 1) a global reduction of NTRK2 during the acute phase in response to hyperoxic injury; 2) increased NTRK2-T1 contributes to impaired vascular recovery and poor outcomes; and 3) NTRK2-FL plays a protective role by promoting alveolar capillary regeneration and repair (**Figure 2O**).

### NTRK2-T1 Exacerbates BPD by Activating RhoA/Calcium Signaling and Impairing ERK/AKT pathways

To elucidate the molecular mechanisms by which NTRK2-T1 contributes to the pathological progression of BPD, we performed pseudo-bulk analysis on gCap ECs from control and BPD patients, with a particular focus on NTRK2-associated signaling pathways. Our analysis revealed that Rho GTPase and calcium signaling pathways were upregulated in BPD gCap cells, whereas PI3K-Akt and MAPK-Ras pathways were suppressed **(Figure 3A)**.

**Figure 3:**
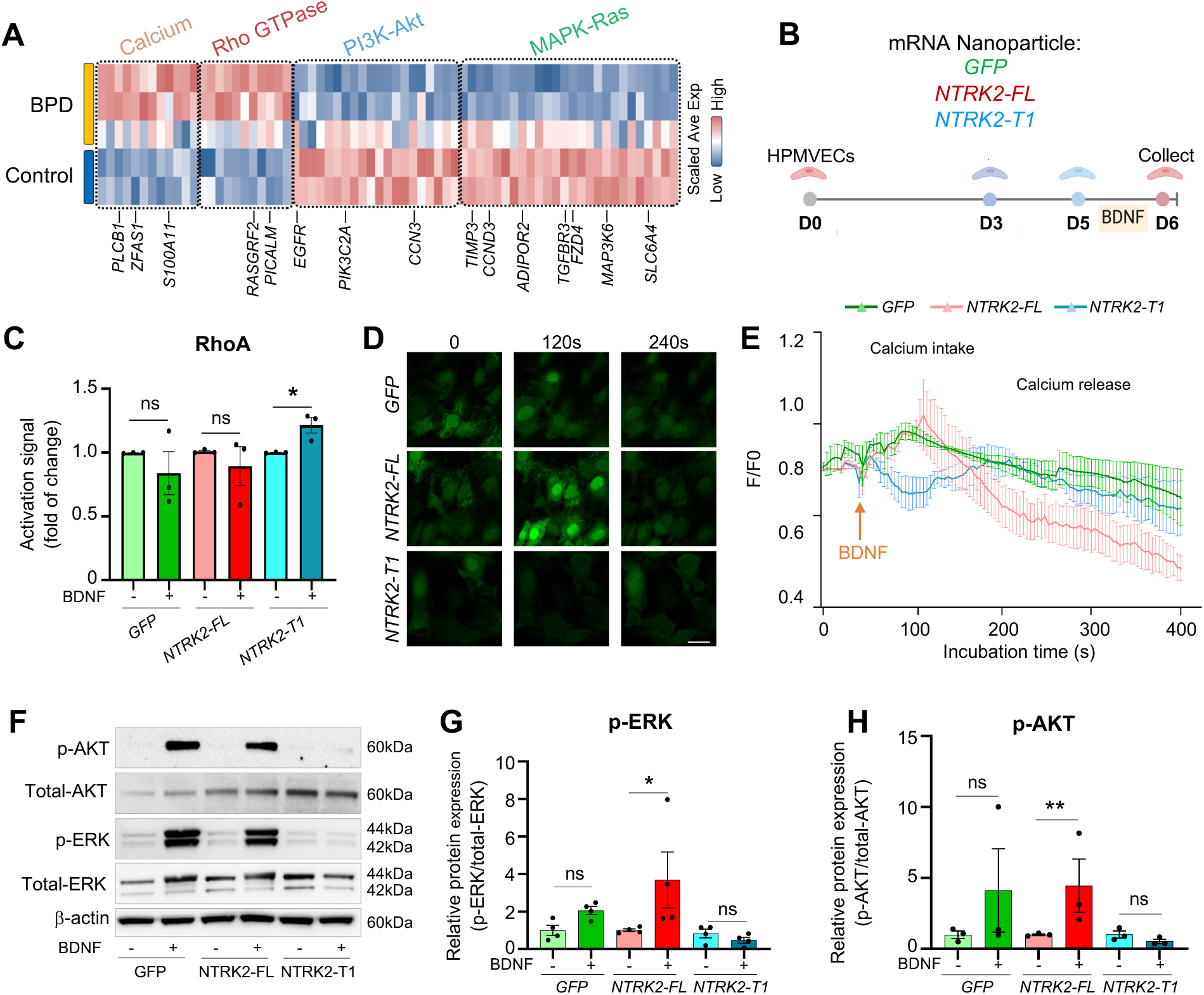
NTRK2-T1 Exacerbates BPD by Activating RhoA/Calcium Signaling and Impairing ERK/AKT pathways (A) Heatmap visualization of expression of Calcium, Rho GTPase, PI3K-Akt, and MAPK-Ras genes differentially expressed in BPD vs. control gCap cells. Each row represents pseudo-bulk gene expression in the gCap cells from the multiome profiling of each sample. Values were per-gene zscore scaled. (B) Experimental design showing that human pulmonary microvascular endothelial cells (HPMVECs) were transfected with either *GFP* mRNA, *NTRK2-FL* mRNA or *NTRK2-T1* mRNA. After transfection for 48 h, the cells were treated with or without BDNF for 24h and then collected. (C) RhoA activation assay in HPMVECs with mRNA transfection and BDNF treatment. Data presented as mean ± SEM, n=3 biological repeats. \**P* < 0.05, unpaired two-tailed *t* test. (D) Representative Fluo-4 Ca²⁺ imaging of HPMVECs transfected with empty vector (*GFP* mRNA), *NTRK2-FL* mRNA, and *NTRK2-T1* mRNA at distinct time points. n = 3 biological repeats. Scale bar, 25 μm. (E) Fluorescent signal quantification of Fluo-4 from (D). Data presented as mean ± SEM. n=3 biological repeats. (F) Western blot of phosphorylation of AKT and ERK in HPMVEC with *NTRK2* isoform mRNA transfection and BDNF treatment. (G) Bar graph of p-ERK/total ERK. Data presented as mean□±□SEM, n=4. \**P* < 0.05, one-way ANOVA followed by Šídák’s multiple comparisons tests. (H) Bar graph of p-AKT/total AKT. Data presented as mean□±□SEM, n=4. \*\**P* < 0.01, unpaired two-tailed *t* test.

To directly assess the impact of NRTK2-FL and NRTK2-T1 signals in pulmonary ECs, we overexpressed either NTRK2-FL or NTRK2-T1 in human pulmonary microvascular endothelial cells (HPMVECs) via nanoparticle mediated mRNA transfection, followed by BDNF stimulation **(Figure 3B, S6A)**. BDNF selectively activated RhoA in NTRK2-T1-overexpressing cells, whereas no significant RhoA activation was observed in NTRK2-FL-expressing or control cells **(Figure 3C)**. Additionally, BDNF-induced transit Ca²⁺ influx was disrupted in the presence of NTRK2-T1, further implicating this isoform in calcium signaling dysregulation **(Figure 3D-3E)**. Moreover, in contrast to NTRK2-FL, overexpression of NTRK2-T1 abolished BDNF-induced phosphorylation of ERK and AKT, key mediators of EC survival and angiogenesis **(Figure 3F-3H)**.

### NTRK2-T1 Lead to Impaired Lung Endothelial Functions

RhoA GTPase, Calcium signaling, and the ERK/AKT pathways are master regulators for cell survival, proliferation, migration, and barrier function. To further investigate NTRK2-T1-mediated vascular dysfunction in BPD, we performed Gene Ontology (GO) analysis of differentially expressed genes (DEGs) in gCap cells from BPD versus control lungs. This analysis revealed significant enrichment of GO terms associated with cell proliferation, angiogenesis, and vascular permeability/barrier function (**Figure 4A**).

**Figure 4:**
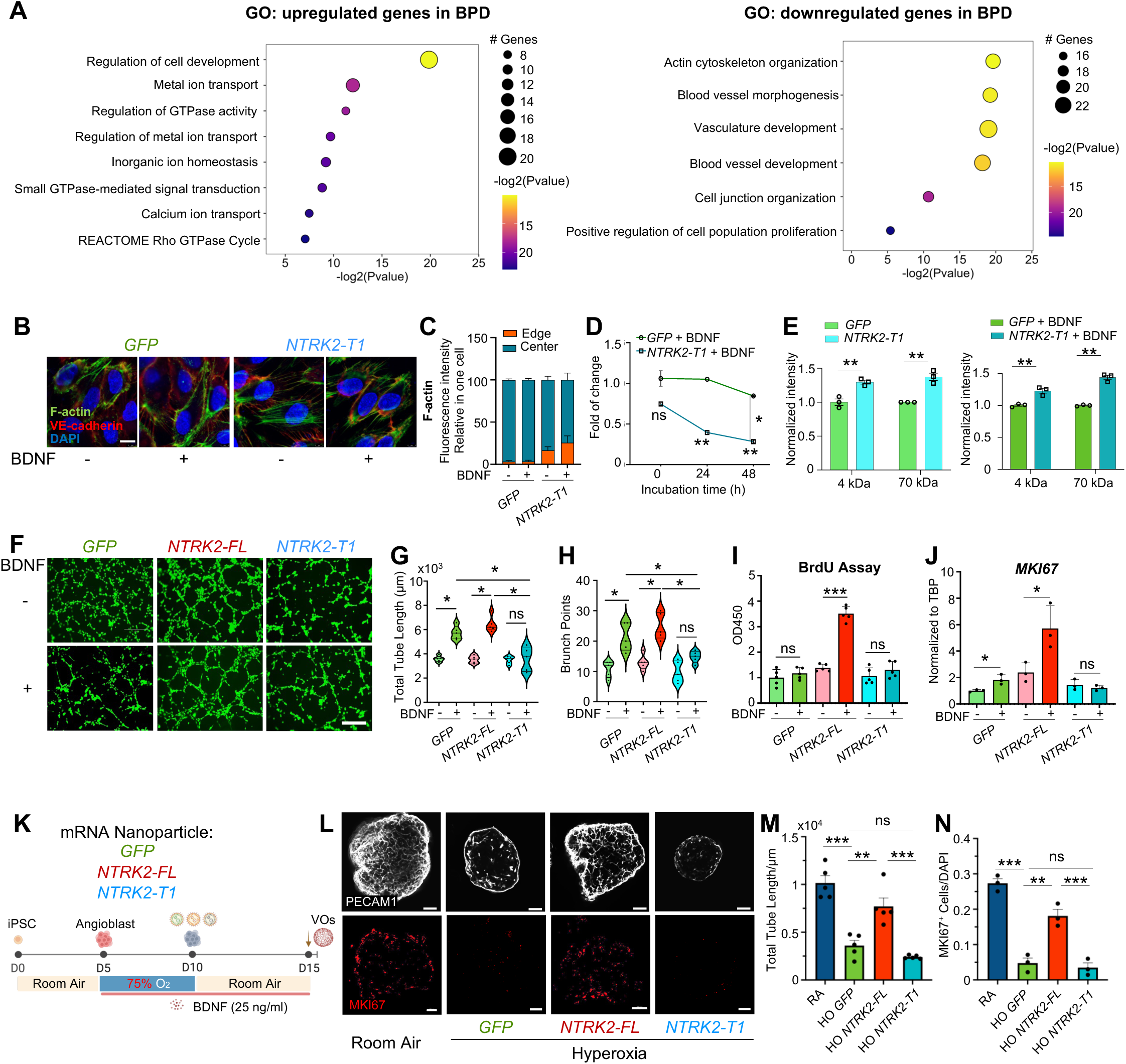
NTRK2-T1 Impairs Vascular Repair in BPD by Disrupting Actin Dynamics, Barrier Function, Proliferation and Angiogenesis *in vitro* (A) Gene ontologies (GOs) enriched by genes differentially expressed in BPD vs. control gCap cells in the multiome profiling. (B) Immunofluorescence staining of F-actin (green) and VE-cadherin(red). Nuclear were stained with DAPI. Scale bar, 5 μm. (C) Quantification of F-actin localization according to panel B was shown in stack bar graph. Data presented as mean ± SEM, n = 6 biological repeats. (D) Trans-endothelial electronic resistance (TEER) of human pulmonary microvascular endothelial cells (HPMVECs) transfected with *GFP* mRNA and *NTRK2-T1* mRNA. Data presented as mean ± SEM, n = 3 biological repeats. *NTRK2-T1*+BDNF at 24h vs. *NTRK2-T1* + BDNF group at 0h, \*\**P* < 0.01; *NTRK2- T1*+BDNF at 48h vs. *NTRK2-T1* + BDNF group at 0h, \*\**P* < 0.01; *NTRK2-T1*+BDNF at 48h vs. *GFP* + BDNF group at 48h, \**P* < 0.05; two-way ANOVA followed by Šídák’s multiple comparisons tests. (E) Bar graph of dextran penetration assay of HPMVEC transfected with *NTRK2-T1* mRNA and treated with BDNF. Data presented as mean ± SEM, n = 3 biological repeats. \*\**P* < 0.01, two-way ANOVA followed by Šídák’s multiple comparisons tests. (F) Angiogenesis assay using HPMVEC with *NTRK2* isoform mRNA transfection and BDNF treatments. Scale bar, 400 μm. (G) Bar graph of total tube length of angiogenesis assay in HPMVECs. Data presented as mean± SEM, n=6 biological repeats. \**P* < 0.05, two-way ANOVA followed by Šídák’s multiple comparisons tests. (H) Bar graph showing brunch points of angiogenesis assay. Data presented as mean± SEM, n=6 biological repeats. \**P* < 0.05, two-way ANOVA followed by Šídák’s multiple comparisons tests. (I) Bar graph showing BrdU signal of HPMVEC transfected with *GFP* mRNA, *NTRK2- FL* mRNA and *NTRK2-T1* mRNA with or without treatment of BDNF. Data presented as mean ± SEM, n = 6 biological repeats. \*\*\**P* < 0.001, two-way ANOVA followed by Šídák’s multiple comparisons tests. (J) Bar graph showing the changes of *MKI67* mRNA via qPCR. Data presented as mean ± SEM, n = 3 biological repeats. \**P* < 0.05, two-way ANOVA followed by Šídák’s multiple comparisons tests, (K) The experimental procedure of vascular organoids (VOs) induced from iPSCs (STAR method). Spheroids were cultured in hyperoxia conditions (75% O₂) from day 5 to day 10. On day 10, *GFP* mRNA, *NTRK2-FL* mRNA, or *NTRK2-T1* mRNA was individually transfected into spheroids. From day 5 to day 15, spheroids were either treated with BDNF or left untreated. (L) Immunofluorescence staining of PECAM-1(white) and MKI67(Red) of day 15 vasculature organoid. Each condition was replicated using 3 iPSC lines. Scale bar:100 μm. (M) Quantification of total tube length in day 15 vessel organoids. Data presented as mean ± SEM; n = 5 vessel organoids. \*\**P* < 0.01, \*\*\**P* < 0.001, one-way ANOVA followed by Bonferroni’s multiple comparisons test. (N) The ratio of MKI67^+^ cells in VOs on day 15. Data presented as mean ± SEM; n = 3 iPSC lines. Each point indicates the average ratio of 5 vessel organoids. \*\**P* < 0.01, \*\*\**P* < 0.001, one-way ANOVA followed by Bonferroni’s multiple comparisons test.

To determine the direct role of NTRK2-T1 in regulating endothelial functions, we overexpressed NTRK2-T1 in HPMVECs and observed profound cytoskeletal disorganization characterized by reduced F-actin accumulation at the cell peripheries (**Figure 4B-4C**), impaired endothelial barrier integrity indicated by decreased trans- endothelial electrical resistance (TEER) (**Figure 4D**), and increased endothelial permeability based on increased dextran flux (**Figure 4E**). Additionally, angiogenic capacity (**Figure 4F-4H**) and cell proliferation (**Figure 4I-4J**) were significantly attenuated in HPMVECs with NTKR2-T1 overexpression, in contrast to the pro- angiogenic effects observed in the NTRK2-FL overexpression group in response to BDNF stimulation (**Figure 4F-J, S6B**).

To further understand the different roles of NTRK2 isoforms in angiogenesis in 3D, we developed a vessel organoid system from human iPSCs. Consistently, NTRK2- T1-overexpressing organoids displayed impaired angiogenic sprouting and reduced proliferative activity. Conversely, organoids overexpressing NTRK2-FL mRNA exhibited robust vascular regeneration and BDNF-responsive angiogenesis under hyperoxic injury (**Figure 4K-4N**), suggesting a potential therapeutic role for NTRK2- FL in mitigating hyperoxia-induced vascular injury.

### Dual Regulatory Axis of HOXA5 and RBFOX2 Contributes to NTRK2-T1 upregulation in BPD

To further understand the mechanism underlying NTRK2-T1 upregulation in BPD, we analyzed the multiomic data from control and BPD gCap cells and identified HOXA5 and MEIS1 binding motifs within the *NTRK2* promoter region which has increased accessibility in BPD gCap cells compared to controls (**Figure 5A**). The expression levels of *HOXA5* was also increased in BPD gCap cells, whereas *MEIS1* levels remained unchanged (**Figure 5B-C, S7A-B**). This was further confirmed by immunofluorescence staining of HOXA5 in lung ECs during both acute and chronic BPD phases (**Figure 5D**) and in hyperoxia-exposed vessel organoids (**Figure S7C**). Gain- and loss-of-function experiments further demonstrated the direct regulation of *HOXA5* on *NTRK2* expression in HPMVECs (**Figure 5E, S7D-E**).

**Figure 5:**
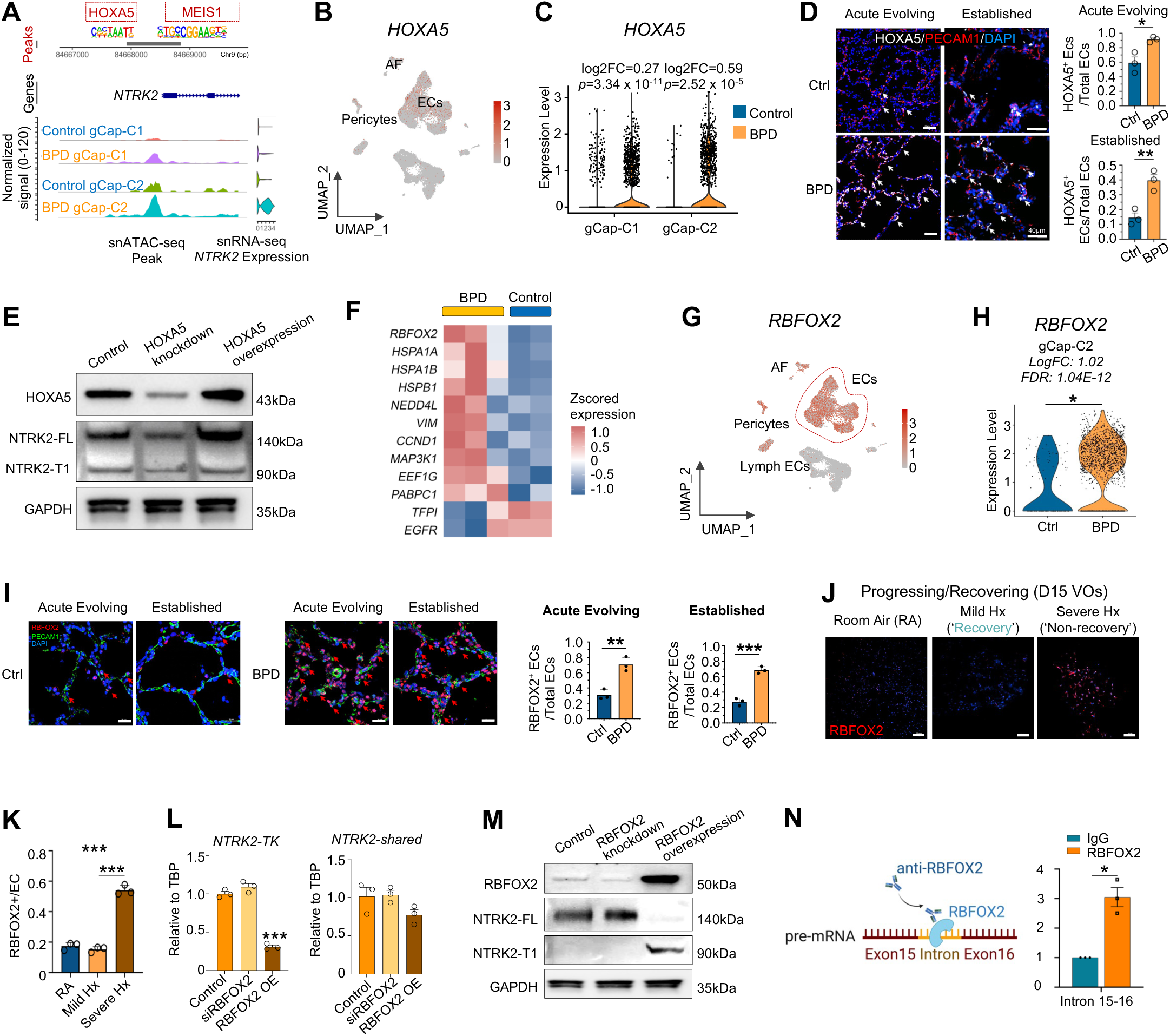
HOXA5-RBFOX2 Synergistically Controlled the Splicing of *NTRK2-T1* **Isoform under Bronchopulmonary dysplasia (BPD) Condition** (A) Prediction of upstream regulators of *NTRK2* using the multiome profiling. HOXA5 and MEIS1 binding sites were predicted in the promoter region of *NTRK2*. (B) UMAP visualization of the expression of *HOXA5* in the multiome profiling of BPD and control lung samples as shown in Figure 1B. (C) Violin plot showing the expression of *HOXA5* in BPD vs. control gCap-C1 or gCap-C2 cells in the multiome profiling. P values and fold change (FC) values were obtained from tests of gene expression in BPD vs. control cells within each cell type using Seurat FindMarkers function using two-tailed Wilcoxon rank sum test. (D) Immunofluorescence image (left) showing HOXA5 expression in distal lung at different stages of BPD. HOXA5(white), PECAM1(red) and DAPI (blue). Scale bar, 40 µm. Bar plot (right) shows the ratio of HOXA5^+^ endothelial cells (ECs) in BPD and control groups during acute evolving stage and established stage. Data represented as mean ± SEM, n = 3 patients. \**P* < 0.05, \*\**P* < 0.01. BPD vs. control, unpaired two-tailed *t* test. (E) Western Blot showing the protein expression of NTRK2 after the knockdown or overexpression of HOXA5. (F) Heatmap visualization of RNA alternative splicing related genes differentially expressed in BPD vs. control gCap cells. Each column represents pseudo-bulk gene expression in the gCap cells from the multiome profiling of each sample. (G) UMAP visualization of the expression of *RBFOX2* in the multiome profiling as shown in Figure 1B. (H) Violin plot showing the expression of *RBFOX2* in BPD vs. control gCap-C2 cells in the multiome profiling. *P* values and fold change (FC) values were obtained from test of gene expression in BPD vs. control gCap-C2 cells using Seurat FindMarkers function using two-tailed Wilcoxon rank sum test. (I) Immunofluorescence images (left) and quantification (right) showing RBFOX2 protein expression in distal lung at different stages of BPD. RBFOX2(Red), PECAM1(Green) and DAPI (blue). Scale bar, 20 µm. Data presented as mean ± SEM, n = 3 patients. \*\**P* < 0.01, \*\*\**P* < 0.001, BPD vs. control, unpaired two-tailed *t* test. (J) Immunofluorescence image showing RBFOX2 expression in vessel organoids in different hyperoxic conditions according to Figure 2J. RBFOX2(Red). Scale bar, 100 µm. (K) Quantification of RBFOX2^+^ ECs in (J). Data represented as mean ± SEM. *n* = 3 iPSC line. \*\*\**P* < 0.001, one-way ANOVA followed by Bonferroni’s multiple comparisons test. (L) qPCR analysis of *NTRK2-TK* region and *NTRK2-shared* region with RBFOX2 knockdown or overexpression in human pulmonary microvascular endothelial cells (HPMVECs). The result was normalized to control and represented as mean ± SEM. n=3. \*\*\**P* < 0.001 vs. Control, one-way ANOVA followed by Bonferroni’s multiple comparisons test. (M) Western Blot showing the protein expression of NTRK2 isoforms induced by the knockdown or overexpression of RBFOX2 in human pulmonary microvascular endothelial cells. (N) The relative pre-mRNA expression of *NTRK2* intron 15-16 after pull-down using RBFOX2 antibody and IgG. Data presented as mean ± SEM, n = 3 repeats. RBFOX2 vs. IgG, \**P* < 0.05, unpaired two-tailed *t* test.

To elucidate the molecular basis of increased alternative splicing of NTRK2 in BPD, we identified RNA-binding proteins (RBP) that differentially expressed in BPD versus controls. Out of the 12 candidate RNA regulators, *RBFOX2* was identified as the most possible RBP driving *NTRK2 FL-*to-*T1* splicing reprogramming (**Figure 5F-H, S7F**).^30^ Elevated *RBFOX2* expression was validated in BPD lung endothelium across disease stages (**Figure 5I**) and in vessel organoids exposed to severe hyperoxic injury (**Figure 5J-K**). *RBFOX2* overexpression in HPMVECs markedly increased *NTRK2-T1* expression and T1/FL ratio (**Figure 5L-M, S7G**). RNA immunoprecipitation (RIP) assay confirmed *RBFOX2* binding to *NTRK2* pre-mRNA at intron 15-16, a splicing binding site for producing *NTRK2-T1* (**Figure 5N**).

Collectively, our findings establish a dual regulatory axis: HOXA5 transcriptionally activates *NTRK2*, while RBFOX2 promotes its splicing toward the truncated isoform, driving vascular dysfunction in BPD.

### Ntrk2-FL RNA Therapy Restores Alveologenesis in BPD-like Mice

To evaluate the therapeutic potential of NTRK2-FL in mitigating hyperoxia-induced alveolar simplification, a hallmark of BPD, we engineered a lipid nanoparticle (LNP) system for the targeted delivery of Ntrk2-FL mRNA specifically to lung ECs via retro- orbital injection (**Figure 6A**, **S8A**). Using *in vivo* luciferase imaging and flow cytometry analysis of dissociated lung cells from Ai14 mice,^31^ we confirmed that the nanoparticles specifically targeting the lung, and preferentially enriched in ECs (**Figure 6B-C**).

**Figure 6:**
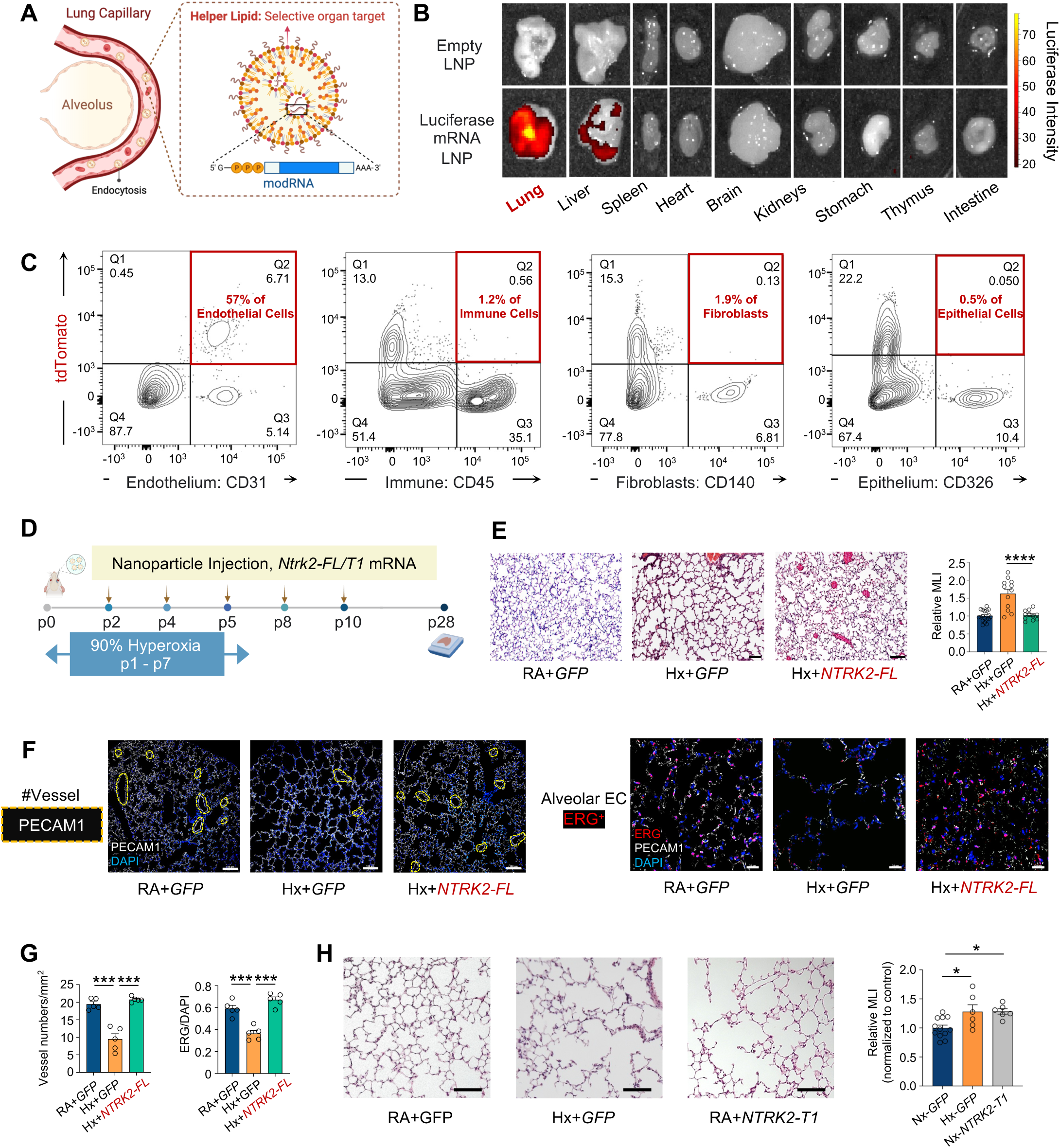
LNP Delivery of *Ntrk2-FL* mRNA Restored Impaired Alveologenesis in Mice Exposed to Hyperoxia Injury. (A) Schematic illustration of the lipid nanoparticle (LNP)-mRNA complex. (B) The luciferase signal in different mouse organs with intravenous (i.v.) administration of luciferase mRNA-LNPs for 48 h (see STAR Methods) *in vivo*. (C) Flow cytometry analysis of dissociated lung cells from Ai14 mice with intravenous (i.v.) administration of Cre mRNA-LNPs for 48 h (see STAR Methods) *in vivo.* % of tdTomato positive cells indicated the uptake efficiency of LNPs in mouse lungs. (D) Experimental timeline for hyperoxic exposure and mRNA-LNP administration (see STAR Methods). (E) Representative H&E staining of distal lungs from postnatal day 28 (P28) mice (from D). Scale bar, 100 μm. Bar graph (right) showing mean linear intercept (MLI) measured by ImageJ. Each point indicates an average of 6 fields per lung section, n = 18 mice from RA + *GFP* group, n = 12 mice from Hx + *GFP* group, n = 11 mice from Hx + *NTRK2-FL* group. Data presented as mean ± SEM. Hx+*NTRK2-FL* vs. Hx+*GFP*, \*\*\*\**P* < 0.0001, one-way ANOVA followed by Bonferroni’s multiple comparisons test. (F) Immunofluorescence staining of distal lung distal lungs from postnatal day 28 (P28) mice (from D). Left: Vessels were labeled by PECAM1(white) and nuclei labeled by DAPI (blue). Scale bar, 100 μm. Right: Alveolar ECs were labeled by ERG (red) and PECAM1(white), nuclei labeled by DAPI (blue). Scale bar, 20 μm. (G) Quantificaition of vessel density and Alveolar EC density. Data presented as mean ± SEM. n = 5 mice. \*\*\**P* < 0.001, one-way ANOVA followed by Bonferroni’s multiple comparisons test. (H) Representative H&E staining images of distal lungs from postnatal day 28 (P28) mice (from D). Scale bar, 100 µm. Bar graph (right) shows MLI of all groups at P28. n = 12 mice for RA + *GFP* group, n=6 mice for Hx + *GFP* group, n= 6 mice for RA + *NTRK2-T1* group. Data are represented as mean ± SEM. \**P* < 0.05, one-way ANOVA followed by Bonferroni’s multiple comparisons test.

Intriguingly, intermittent administration of LNPs containing Ntrk2-FL mRNA significantly mitigated hyperoxia-induced impairments in angiogenesis and alveolar development in murine BPD models (**Figure 6D-G**). In contrast, pharmacological activation of NTRK2 with small-molecule agonist 7,8-DHF failed to elicit any therapeutic benefits, nor does antagonist ANA-12 cause more severe damage (**Figure S8B-C**), indicating that BPD pathology stems from receptor dysfunction, rendering ligand-based activation ineffective.

To further elucidate the detrimental role of *NTRK2-T1* in neonatal lung development, we conducted a parallel experiment in which NTRK2-T1 mRNA nanoparticles were administered to newborn mice under normoxic conditions. Notably, overexpression of *Ntrk2-T1* in neonatal mice resulted in impaired alveologenesis during postnatal lung development (**Figure 6H**), supporting its role in causing lung endothelial dysfunction.

These findings suggest that NTRK2-FL RNA therapy represents a promising therapeutic strategy for BPD, offering potential avenues for restoring vascular and alveolar development in affected neonates.

To determine whether the role of NTRK2-FL in vascular regeneration is lung specific, we evaluated the effect of Ntrk2 in a hindlimb ischemia model by inducing post- ischemic injury in BALB/c mice. Our results demonstrated that Ntrk2-FL-LNP treatment significantly improved blood flow compared to the control group without significant body weight loss observed in either group throughout the experimental period (**Figure S8D-E**). immunofluorescence staining for CD31 revealed a marked increase in vascular density in the Ntrk2-FL-treated group compared to controls (**Figure S8F-G**), supporting our observation from doppler imaging.

## DISCUSSION

In this study, we identify an NTRK2 isoform switch as a key pathogenic driver of alveolar capillary dysfunction in hyperoxia-induced lung injury (**Figure 7**). By integrating multiomic profiling, spatial transcriptomics, and functional validation, we demonstrate that the balance between NTRK2-FL and NTRK2-T1 governs the regenerative capacity of alveolar capillaries. These findings uncover a previously unrecognized therapeutic axis in BPD, a disease with limited treatment options.

**Figure 7:**
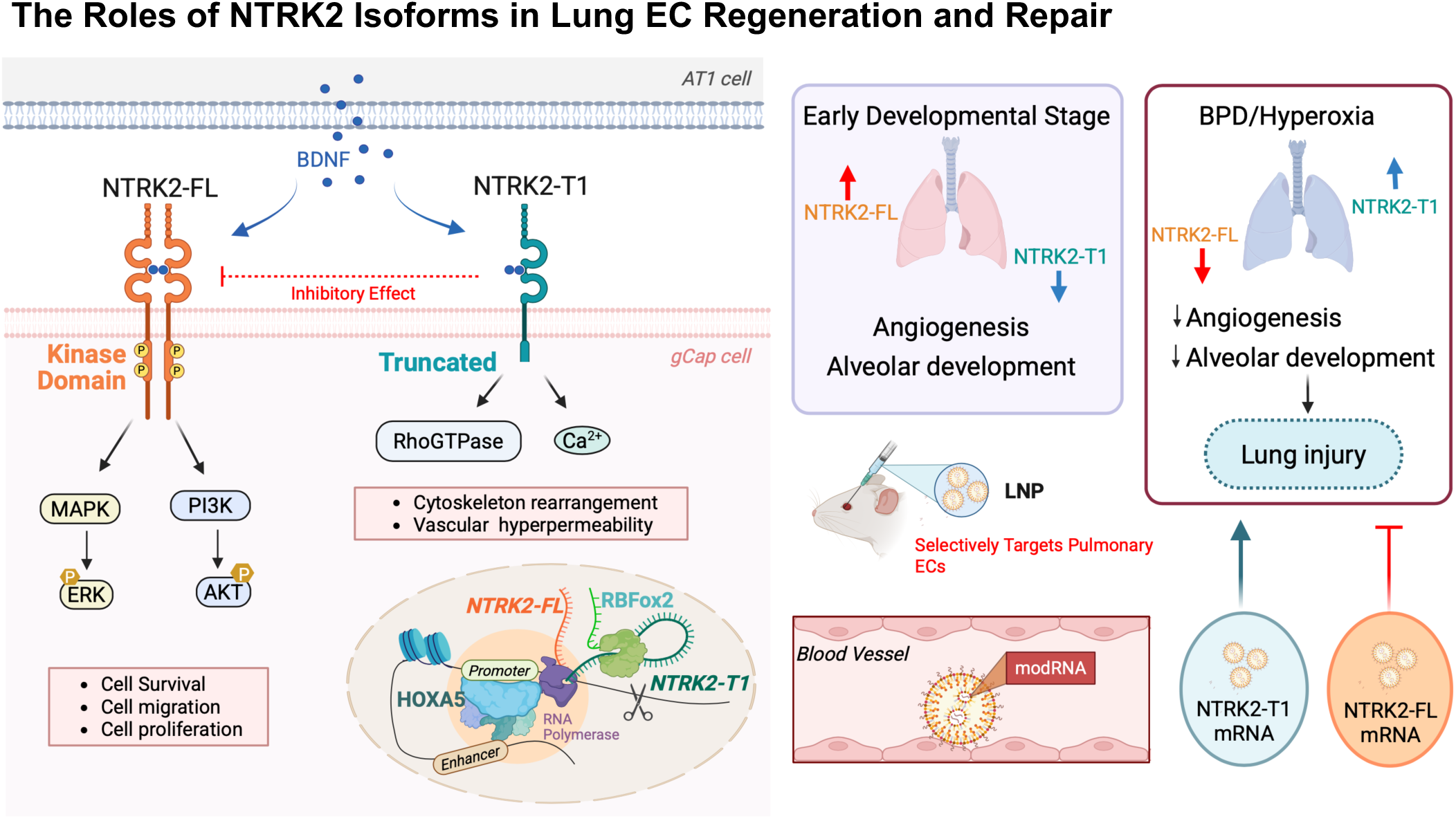
Regulatory Mechanism of Different NTRK2 Isoforms in Lung ECs. Schematic overview of the differential signaling and developmental roles of NTRK2 isoforms in pulmonary ECs. Left panel: NTRK2-FL, possessing an intracellular kinase domain, activates MAPK/ERK and PI3K/AKT pathways to promote endothelial cell survival, migration, and proliferation. In contrast, NTRK2-T1 lacks the kinase domain and engages RhoGTPase and Ca²⁺ signaling, leading to cytoskeletal rearrangement and vascular hyperpermeability. Transcriptional regulation of isoform expression involves HOXA5 induced NTRK2 expression and RBFOX2 mediated splicing of NTRK2, resulting in increased NTRK2-T1 expression. Right panel: During early lung development, NTRK2-FL is upregulated to support angiogenesis and alveolarization, while NTRK2-T1 remains low. In BPD or hyperoxic conditions, this balance is disrupted, with reduced NTRK2-FL and elevated NTRK2-T1 leading to impaired vascular and alveolar development and irreversible lung injury. Lipid nanoparticle (LNP)-mediated delivery of *NTRK2-FL* mRNA selectively targeting pulmonary ECs restores alveolar development in BPD-like mice.s

General capillary endothelial (gCap) cells, located primarily within the thin-walled alveolar septa, play a central role in maintaining vascular integrity and facilitating gas exchange.^32,33^ Using human lung tissues from control and BPD cases, we identified a disease-specific expansion of the gCap-C2 endothelial subpopulation, characterized by its localization to regions of alveolar simplification and a low proliferative profile. To further understand how aberrant gCap-C2 cells affect surrounding tissues, we performed spatial transcriptomic analysis on human BPD lung sections and identified abnormal ligand–receptor interactions between gCap-C2 cells and adjacent fibroblasts, including *EDN1–EDNRB* signaling, indicating disrupted endothelial– mesenchymal communication. Disruption of these signaling axes by hyperoxia may impair paracrine signaling, leading to weakened vascular integrity and impaired alveolar regeneration, which together contribute to the persistent simplification of the alveolar–capillary unit in severe BPD.

While *NTRK2* is a well-established regulator in the central nervous system, where it modulates vascular integrity and tone via BDNF signaling,^34–38^ its role in the pulmonary endothelium has only recently been recognized in mouse models. Injury-induced capillary endothelial cells (iCAPs) with high *Ntrk2* expression and aberrant regenerative traits have been identified in chronic lung injury,^20^ and *Ntrk2*⁺ capillary ECs have been shown to persist near fibrotic lesions in aged mice following bleomycin-induced injury.^21^ However, these studies did not examine the specific roles of individual *NTRK2* isoforms, which we identified as key regulators of alveolar capillary regeneration in humans. In our work, we systematically evaluated the dynamic changes of different *NTRK2* isoforms during alveologenesis and in response to mild versus severe hyperoxic injury at both acute and chronic stages, providing a mechanistic basis for isoform-specific therapeutic strategies in BPD.

Furthermore, we identified HOXA5 and RBFOX2 as key upstream regulators of *NTRK2* isoform imbalance, suggesting a link between transcriptional and post- transcriptional modification of endothelial repair. HOXA5, a transcription factor essential for lung morphogenesis, has been implicated in pulmonary fibrosis and lung cancer, where it inhibits angiogenesis by suppressing VEGF signaling.^39,40^ In BPD, we observed HOXA5 upregulation in gCap cells, contributing to the overall upregulation of NTRK2 in response to hyperoxic injury. Beyond transcriptional regulation, we identify RBFOX2 as a central splicing factor driving the shift from *NTRK2-FL* to *NTRK2-T1* under sereve, non-recovery hyperoxic stress. RBFOX2 is an RNA-binding protein regulating alternative splicing programs in cardiovascular and neurological systems, impacting processes such as cardiomyocyte maturation and vascular remodeling.^41–43^ Our study reveals a previously unrecognized role for RBFOX2 in lung endothelial cells, where its upregulation under hyperoxic conditions drives alternative splicing of full-length *NTRK2*, thereby impairing vascular regeneration and compromising barrier integrity. These findings are consistent with prior work showing RBFOX2 involvement in diabetic retinopathy and pulmonary arterial hypertension,^44^ positioning it as a conserved regulator of vascular injury responses. Notably, RBFOX2 function may differ depending on its subcellular localization. In our study, nuclear RBFOX2 enhances *NTRK2* splicing, whereas the role of cytoplasmic RBFOX2 in regulating mRNA stability that previously attributed to RBFOX1,^30,45^ remains to be investigated. Understanding the spatial regulation of RBFOX2 and its impact on endothelial fate may unlock new therapeutic strategies to reprogram splicing decisions during vascular injury.

Although previous studies have reported beneficial effects of the NTRK2 agonist 7,8- DHF in acid-induced lung injury models,^46^ our mechanistic investigations revealed that under hyperoxic conditions, the primary defect lies in receptor splicing, where full-length *NTRK2* is converted into a nonfunctional truncated isoform. As a result, receptor activation with an agonist is ineffective. To overcome this, we developed a lipid nanoparticle (LNP) system to deliver *Ntrk2-FL* mRNA directly and specifically to lung endothelial cells. This platform builds upon recent successes of mRNA-LNP therapies, which have transformed clinical care in diseases such as COVID-19 and cystic fibrosis.^47–49^ Unlike traditional treatments aimed at symptomatic relief or nonspecific growth factor stimulation (e.g., VEGF or FGF supplementation), our method targets the core molecular defect, the loss of kinase-active NTRK2 signaling. This precision strategy reversed alveolar simplification and restored capillary density in hyperoxia-induced lung injury mouse models. This result supports the crucial insight that receptor isoform dysregulation rather than ligand deficiency is the principal barrier to vascular regeneration in BPD. We further demonstrated that *NTRK2-FL* mRNA not only promotes vascular regeneration in hyperoxic lung injury but also restores vascular function in a hindlimb ischemia model, where endothelial cells are subjected to hypoxic stress. This finding suggests broader therapeutic potential for modulating *NTRK2* isoform expression in oxygen-related vascular injuries. However, additional studies are needed to validate these results and to explore potential differences across various models of vascular injury.

To evaluate therapeutic efficacy in a human-relevant context, we employed iPSC- derived vessel organoids that recapitulate key features of human endothelial phenotypes and injury responses. Overexpressing *NTRK2-FL* mRNA promotes BDNF-responsive angiogenesis under hyperoxia conditions, which provides a predictive platform for therapy validation in a BPD-mimetic microenvironment.^50^ Taken together, our findings support *NTRK2-FL* mRNA delivered via lipid nanoparticles specifically targeting lung endothelial cells as a promising and translationally relevant therapeutic strategy for neonatal lung disease, validated in both human vessel organoids and relevant mouse models.

In conclusion, our study identifies *NTRK2* isoform dysregulation as a central mechanism driving persistent vascular defects in BPD. By restoring the balance of *NTRK2* isoforms through targeted mRNA delivery of *NTRK2-FL* via lipid nanoparticles, we establish a receptor-specific therapeutic strategy that effectively reestablishes vascular integrity. These findings redefine the molecular basis of neonatal vascular injury and position isoform-specific RNA therapy as a powerful platform for regenerative medicine in the developing lung. Future studies will be critical to explore the broader applicability of this approach in other models of vascular injury and to support its clinical translation for neonatal care.

### Limitations of the study

While our findings provide strong evidence that *NTRK2* isoform imbalance contributes to endothelial dysfunction in BPD, several limitations should be acknowledged. First, although iPSC-derived vessel organoids offer a human-relevant platform, they do not fully replicate the complex multicellular architecture and biomechanical cues of the developing lung. Future integration of multi-lineage, perfused organoid systems or lung-on-chip platforms may more accurately model the dynamic interactions among endothelial, epithelial, and immune cells during injury and repair. Second, while we identify RBFOX2 as a key splicing regulator in response to hyperoxia, the upstream pathways controlling its expression, activity, and subcellular localization remain poorly understood. Investigating the roles of redox signaling, epigenetic regulation, and chromatin remodeling could uncover additional mechanisms that influence *NTRK2* isoform switching and provide novel targets for intervention. Finally, although *NTRK2-FL* mRNA therapy effectively restored vascular integrity in vivo, the durability and temporal dynamics of isoform correction remain uncertain. Given the transient nature of mRNA expression, strategies such as repeated dosing or the use of self-amplifying RNA platforms may be required to sustain therapeutic effects. Addressing these limitations will deepen our understanding of vascular injury and repair in BPD and advance the development of durable, scalable, and clinically translatable isoform-targeted therapies for neonatal lung disease.

## Supporting information

Supplemental images

## RESOURCE AVAILABILITY

Lead contact

Further information and requests for resources and reagents should be directed to and will be fulfilled by the lead contact, Mingxia Gu.

Materials availability

This study did not generate new, unique reagents.

## Data and code availability

- Processed 10x single nucleus multiome and 10x Visium spatial transcriptomics data is accessible through the project github: https://github.com/MZGuo-lab/NTRK2_BPD. We will submit the raw and processed sequencing data to dbGaP and GEO, respectively.
- Original western blot images will be deposited at Mendeley and are publicly available as of the date of publication. Microscopy data reported in this paper will be shared by the lead contact upon request.
- R code used in the single cell and spatial analysis is available at: https://github.com/MZGuo-lab/NTRK2_BPD.
- Any additional information required to reanalyze the data reported in this paper is available from the lead contact upon request.

## AUTHOR CONTRIBUTIONS

C.T., Z.L., Y.M., and M.Gu designed the project. C.T, Z.L., Y.M., X.M., N.P., and X.L. performed experiments, supported data analysis, and assisted with interpretation of results. Z.L., and M.Guo. performed computational analysis. H.F., Y.L., contributed to sample preparation for spatial transcriptomics. X.L. generated lipid nanoparticles. V.K., assisted in establishing BPD mouse models. G.P. provided BPD human tissues. R.L., and L.L., performed experiments using hindlimb ischemia mouse model. C.T., X.M., Y.Z., Y.M., and M.Gu wrote the manuscript. All authors reviewed, provided feedback, and approved the manuscript.

## ACKNOWLEDGMENTS

We gratefully acknowledge the Discover Together BioBank for their foundational support of this study. We extend our deepest gratitude to all participants and their families, whose invaluable contributions made this research possible. We thank Dr. Matt Kofron from the Cincinnati Children’s Bio-Imaging and Analysis Facility and Joseph Kitzmiller (Division of Pulmonary Biology, Cincinnati Children’s Hospital Medical Center, CCHMC) for their technical expertise in confocal microscopy and image processing. We thank Drs. Jeffrey A. Whitsett, Kathryn A. Wikenheiser- Brokamp, Anne Karina Perl, and Matthew Riccetti for sharing lung tissue blocks of the BPD mice and guidance on interpreting human BPD pathology. Special thanks to the Research Flow Cytometry Facility at CCHMC for their exceptional assistance in fluorescence-activated cell sorting (FACS) and data analysis. This work was supported by the CCRF Endowed Scholar Award (M.Gu) and the NIH/NHLBI R01HL166283 (M.Gu) and N.P. was funded by the American Heart Association Pre- Doctoral Fellowship (Grant #1013861).

## SUPPLEMENTAL INFORMATION

**Document S1.** Supplemental Figures S1–S8 (this is the main PDF)

## STARfllMETHODS

### KEY RESOURCES TABLE

SEE TABLE 5

### EXPERIMENTAL MODEL AND STUDY PARTICIPANT DETAILS

#### Human Lung Samples

Human lung samples were obtained from infants diagnosed with bronchopulmonary dysplasia (BPD) (ages 8–21 months; n=3) and age-matched controls (n=2) from the Biorepository for Investigation of Neonatal Diseases of the Lung (BRINDL) (Table 1). Tissues were enzymatically digested to generate single-cell suspensions according to previous reported method.^22^ ECs were enriched by magnetic-activated cell sorting (MACS) using CD31^+^/CD45^-^ markers, with cell purity validated by gene count, molecular count, and mitochondrial DNA content criteria. Human lung sample blocks were obtained from BRINDL. Patient information: Table 4.

#### Animal Model of Hyperoxia-Induced Lung Injury

Newborn WT C57BL/6J mice (breeding pairs were obtained from The Jackson Laboratory) were placed in either RA or into a hyperoxia O_2_ propylene chamber (30″

× 20″ × 20″) in which the oxygen concentration was maintained at 75 % or 90 % O_2_, according to previous report .^51^ Neonatal mice were exposed to hyperoxia condition from PN0–PN7 then returned to RA until being harvested at PN10 or PN28. Nursing dams were rotated between groups every 24 h to prevent oxygen toxicity damage to the dams. All mice were maintained in 12/12□h light/dark cycle and received food and water ad libitum. All animal procedures were approved by the Animal Care Committee of the Cincinnati Children’s Hospital (protocol IACUC 2022-0082).

#### Murine hindlimb ischemia model

Male mice were used in the hindlimb ischemia model as previously described.^51,52^ Briefly, the proximal and distal femoral arteries were ligated. Blood flow to the ischemic and unoperated hindlimbs was quantified using a PeriScan PSI NR Laser Doppler system (Perimed AB, Sweden) as described previously. Limb perfusion was monitored immediately after surgery, and 3, 7, 14, 21 and 28 days postoperatively. Limb perfusion was measured via laser Doppler scans over a 30-second cycle at a frame rate of 6 images/second. The region of interest (ROI) was the hindlimb distal to the knee, including the foot, which accounts for the majority of hindlimb perfusion and is a clinically relevant area of interest. The level of perfusion in ischemic and unoperated hindlimbs was quantified via mean pixel value within the ROI. To account for variation in temperature, ambient light, and arterial pressure, the relative change in hindlimb perfusion was expressed as a perfusion index, the ratio of ischemic over nonischemic (contralateral) laser doppler detected blood perfusion. Mice were weighed at the time of surgery, and on postoperative days 3, 7, 14, 21 and 28, to make sure there are no weight differences between treatment groups during recovery.

#### Lipid nanoparticle-encapsulated mRNA (LNP-mRNA) preparation

The mRNA-loaded LNP formulation was prepared using the ethanol dilution method .^53^ Specifically, lipids were dissolved in ethanol with following concentrations: Ckk-E12 (10 mg/mL), 18:0 DDAB Dimethyldioctadecylammonium bromide Salt (18:0

DDAB) (5 mg/mL),1,2-dimyristoyl-sn-glycero-3-phosphoethanolamine-N- [methoxy(polyethyleneglycol)-2000](ammonium salt)(C14-PEG2000) (5 mg/mL), and cholesterol (5 mg/mL). These solutions were combined at a volume ratio of 35:44.5:2.5:18 to form the organic phase.

mRNA was dissolved in 20 mM citrate buffer (pH 3.0), and the organic phase added dropwise into the aqueous phase at a 1:3 volume ratio using a 1ml syringe. The mixture was allowed to react for 5 min. To remove ethanol and adjust the pH, PBS (pH 7.4) was added, and the solution was transferred to an Amicon Ultra centrifugal filter (3 kDa MWCO) for centrifugation. The washing step was repeated three times, yielding a final pH of approximately 7.

The final solution was filtered through a 0.22 µm filter and prepared for in vivo experiments. The fresh LNP formulation was diluted with PBS for size detection using dynamic light scattering (DLS).

#### Cell Lines

##### Human Induced Pluripotent Stem Cells (iPSCs)

All human iPSC work was approved by Institutional Review Boards at Cincinnati Children’s Hospital, Boston University, and Erasmus University Medical Center Rotterdam. Three iPSC lines used included: control-1 (BU3 NGST, male, 32 years), control-2 (CIRM iPSC Repository, female, 1 year), and control-3 (BRINDL-derived, female, 20 months). iPSCs were maintained in StemMACS iPSC-brew XF medium (Miltenyi Biotec) on Cultrex (Bio-Techne)-coated 6-well plates at 37 °C, passaged every 3–5 days, and authenticated routinely for karyotype, pluripotency markers, STR analysis, and mycoplasma contamination.

#### Primary Human Pulmonary Microvascular Endothelial Cells (HPMVECs)

HPMVECs (Lonza) were cultured in endothelial cell growth medium-2 (Lonza) supplemented with fetal bovine serum (5%), growth factors (hEGF, VEGF, bFGF, IGF-1), ascorbic acid, hydrocortisone, and antibiotics. Cells were maintained on 0.1% Type I collagen-coated flasks (Advanced Biomatrix) at 37°C in a humidified atmosphere containing 5% CO₂ until grown to 80–90% confluence and passaged using trypsin-EDTA (Fisher Scientific). Medium was refreshed every 48 h. Cell viability (>95%) was confirmed by trypan blue exclusion before experimental use.

### METHOD DETAILS

#### LNP-mRNA treatment in Animal Model of Hyperoxia-Induced Lung Injury

Animal models of hyperoxia-induced lung injury were prepared as previously described. Briefly, newborn pups were continuously exposed to 90% oxygen in sealed chambers from P0 to P7, followed by maintenance in room air until tissue collection at P28. LNP-mRNA complexes containing therapeutic mRNA were administered via transorbital injection at a dose of 1 μg mRNA per gram body weight (10 μL/g) on postnatal days 2, 4, 6, 8, and 10 (Hx + NTRK2-FL mRNA group). Hx + GFP mRNA group received equivalent doses of GFP-encoding LNP-mRNA complexes through the same administration route. RA + GFP mRNA group received equivalent doses of GFP-encoding LNP-mRNA complexes without hyperoxia treatment.

#### Ntrk-T1 LNP-mRNA induced lung injury model

Newborn pups were raised in room air from P0 to P28. *Ntrk2-T1* LNP-mRNA complexes were administered via transorbital injection at a dose of 1 μg mRNA per gram body weight (10 μL/g) on postnatal days 2, 4, 6, 8, and 10. RA + GFP mRNA group and Hx + GFP mRNA group were generated as described above.

#### NTRK2 agonist and antagonist treating hyperoxia induced lung injury model

Animal models of hyperoxia-induced lung injury were prepared as previously described. Briefly, newborn pups were continuously exposed to 90% oxygen in sealed chambers from P0 to P7, followed by maintenance in room air until tissue collection at P28. Pups received intraperitoneal injections of 10□μg/g 7,8-DHF (Sigma–Aldrich) or 1 μg/g ANA-12(Selleckchem) at P8, 10, 12, 14, 16. Hx + DMSO and RA + DMSO group received equivalent doses DMSO with or without hyperoxia treatment.

#### Lung isolation

Pups were euthanized at P28 by injection of 50 uL of triple sedative solution (67 mg/mL ketamine + 3.3 mg/mL xylazine + 1.7 mg/mL acepromazine) by intraperitoneal injection and a method of secondary euthanasia was employed (exsanguination by severing the carotid artery). Adult mice used in the organoid study were euthanized in the same manner but received 100 ul of triple sedative solution. Following euthanasia, mice were tracheotomized and lungs were installation-fixed for 5□min at 25□cm H2O hydrostatic pressure with 4 % (w/v) paraformaldehyde (PFA) fixation solution (Fisher Scientific). After isolation, lungs were kept in the fixation solution for 48□h at 4□°C and collected for embedding in paraffin. Paraffin-embedded tissue blocks were sectioned at 4 μm and stained with hematoxylin and eosin (H & E) stain.

#### Isolation of gCap cells in mouse lungs

Lungs were removed from ice-cold PBS and non-pulmonary tissue and gross airways were removed via manual dissection, and lung tissue was finely chopped and transferred to a GentleMACS C tube (Miltenyi Biotec) (tissue from one mouse per C tube) containing 5□mL of digestion buffer [composed of 9□mL of phosphate-buffered saline (PBS; Gibco) combined with 1□mL of Dispase (stock: 50□U/mL; final concentration: 5□U/mL, Corning), 50□µL of DNase (stock: 5□mg/mL; final concentration: 0.025□mg/mL or 50 U/mL, GoldBio), and 100□µL of Collagenase Type I (stock: 48,000□U/mL; final concentration of 480□U/mL, Gibco)]. C tubes were placed on a gentleMACS Octo Dissociator with Heaters (Miltenyi Biotec), and the following protocols were run: “m_lung_01_02” (36□s) twice, “37C_m_LIDK_1” (36□min 12□s) once, and “m_lung_01_02” (36□s) once. Samples were passed through a 70□µm filter (Greiner Bio-One) and centrifuged at 500□g for 5□min at 4□°C. Following removal of the supernatant, 5□mL of RBC Lysis Buffer (Invitrogen) was added and incubated for 5□min. All centrifugation steps with this single-cell suspension were performed at 500 g for 5□min at 4□°C for the following procedures.

After RBC lysis, the single-cell suspensions were washed twice with cold Cell Staining Buffer (BioLegend) and adjusted to 1 × 10□ cells/mL in DPBS containing 0.1 % BSA. Added 1:1000 dilution of Fixable Viability Dye BV510 (BD Biosciences) and Incubated 20 min at 4 °C in dark with gentle vertexing every 5 min. the reaction was quenched by adding 5 × volume of cold cell staining buffer and washed by cold DPBS for twice. Then the cell was stained by c-Kit-PE (1:100, Biolegend) and Pecam1-APC (1:100, Biolegend) for 20 min at 4 °C in dark. Then the cells were washed with cell staining buffer for twice. Kit^+^/Pecam1^+^ cells were sorted using BD FACSAria Fusion cell sorter.

#### iPSC-derived Vessel Organoids (VOs)

Human iPSC-derived VOs were generated based on a published method,^54^ with some slight modifications. hiPSCs were first dissociated into single cells and resuspended in Aggregation media consisting of Knockout DMEM/F12 (Gibco), 20% Knockout Serum Replacement (Gibco), 1 % GlutaMAX™, 1 % MEM Non-essential Amino Acid Solution (Gibco), 55 μM beta-mercaptoethanol (Gibco), and 1 % Anti-Anti (Gibco) supplemented with 20 µM Y-27632 (Tocris). 1.2 × 10^6^ cells were added to each well of the Aggrewell™400 (Stem Cell Technologies) that was already prepped with Anti-Adherence Rinsing Solution (Stem Cell Technologies) per manufacturer’s instructions. The Aggrewell™400 consisting of cells was then spun down at 100 g for 3 min to evenly distribute cells into each microwell (1000 cells per microwell). Cells were then left to form embryoid bodies (EBs) overnight at 37 °C. The next day, EBs were transferred to N2B27 media supplemented with 12 μM CHIR99021 (Selleck Chem) and 30 ng/mL BMP4 (R&D Systems) and cultured in ultra-low attachment 6 well plates (Corning) on an orbital shaker at 37 °C for three days. Subsequently, the media was changed to N2B27 supplemented with 2 μM Forskolin (Sigma-Aldrich) and 100ng/ml VEGFA-165 (GeminiBio) for 2 days. Then, on day 5, the aggregates were embedding into a Collagen I-Matrigel sandwich consisting mixture consisting of 11.3 % 0.1 N NaOH (Sigma Aldrich), 4.7 % 10 × DMEM (Thermo Fisher Scientific), 0.9 % 1 M HEPES (GibcoTM), 0.7 % NaHCO_3_ (GibcoTM), 0.5 % GlutaMAX (GibcoTM), 6.9 % Ham’s F-12 (GibcoTM), 50% PureCol® (Advanced BioMatrix), and 25 % Cultrex RGF Basement Membrane Extract Type 2 (R&D Systems). To induce vessel sprouting, blood vessel induction (BVI) media composed of StemPro™-34 SFM media supplemented with StemPro™-34 nutrient mix (Gibco), 1 % GlutaMAX™ (Gibco), 1% Antibiotic-Antimycotic (Gibco), 15 % Fetal Bovine Serum (FBS) (Gibco), 100ng/ml VEGFA-165 (Gemini Bio), and FGF2 (Gemini Bio) was added to the collagen-matrigel sandwiches. Fresh BVI media was changed every 2 days. On day 10, the vessel sprouts were then micro-dissected from the collagen-Matrigel sandwich gels using 27G sterile needles (BD Biosciences) and transferred to ultra-low attachment 6 well plates (Corning) containing fresh BVI media and cultured on an orbital shaker at 37 °C for 24 h to form VOs. Subsequently, the VOs were transferred to ultra-low attachment 96-well plates (S-bio) and supplemented with fresh BVI for the remaining 4 days. BVI media was replaced every 2 days and VOs were harvested on day 15 of differentiation unless otherwise stated. To model oxidative stress, organoids were transferred to hyperoxic conditions (60 % O₂ until Day 8 or 75 % O₂ until Day 10) starting on Day 5, alongside continuous supplementation of 25 ng/mL BDNF until Day 15.

#### Overexpression and RNAi

##### mRNA transfection in HPMVEC and vessel organoid

Primary human pulmonary microvascular endothelial cells (HPMECs) at passages 4-6 or Day 10 blood vessel organoids were transfected using jetMESSENGER transfection reagent (Polyplus) with the following optimized protocol. For HPMVECs, Cells were seeded at 1.5×10□ cells/cm² in type I collagen-coated (Cornin) 6-well plates and cultured in antibiotic-free EGM-2 MV medium (Lonza, CC-3202) for 24 hr to attain 70-80% confluency prior to transfection. For blood vessel organoids, organoids were cultured in 6 well plates at 15-30 organoids per well.

Transfection complex preparation: 1 μg of purified mRNA (*GFP*, *NTRK2-FL or NTRK2-T1*) was diluted in 200 μL sterile jetMESSENGER buffer (provided with 717- 05 kit); Combined with 2 μL jetMESSENGER reagent (1:2 mRNA: reagent mass ratio); Vortexed vigorously for 10 sec followed by 15 min incubation at room temperature (22-25°C); The complex solution was diluted in 2 mL pre-warmed Opti-MEM reduced serum medium (Gibco); The diluted complexes were applied dropwise to cell monolayers with gentle plate agitation. After 4 h incubation under standard culture conditions (37°C, 5% CO₂), the transfection medium was aspirated and replaced with complete EGM-2 MV medium supplemented with 10% FBS (Gibco). Following transfection, HPMEC cultures were subjected to recombinant human BDNF (PeproTech,) treatment at 25 ng/mL in serum-reduced medium (EGM-2 MV containing 2% FBS). The functional assay and signaling pathway activation validation was performed 24h after BDNF treatment.

#### Plasmid and RNAi transfection in HPMVEC

HPMVECs at passages 4-6 were transfected using transfection reagent (Polyplus) with the following optimized protocol. At 70–80% confluency, cell cultures on 6 well plates were briefly washed with warm Dulbecco’s phosphate buffered saline (PBS, Thermo Fisher Scientific) and change to 1ml pre-warmed Opti-MEM reduced serum medium (Gibco). For jetOPTIMUS, 1µg DNA was added into 100 µL of supplied buffer and gently vortexed. To that, 1 µL of jetOPTIMUS was added into the buffer at a reagent: DNA ratio of 1:1. The mixture was gently vortexed and incubated at room temperature for 10 min before added to the well. After 4 h incubation under standard culture conditions (37°C, 5% CO₂), the transfection medium was aspirated and change to RNAi transfection complex. For RNAi transfection complex, siRNA-SMARTpool (Dharmacon) and Lipofectamine RNAiMAX (Thermo Fisher Scientific) were complexed in Opti-MEM (Gibco) as follows: 20 nM siRNA (final concentration) was diluted in 25 μL Opti-MEM, while Lipofectamine RNAiMAX was diluted at a 1:100 ratio (v/v) in a separate 25 μL Opti-MEM. The diluted siRNA and Lipofectamine RNAiMAX were combined, gently mixed, and incubated at room temperature for 10 min to form siRNA-lipid complexes. The complexes (50 μL total volume) were then added dropwise to each well containing HPMVECs in serum-containing medium. Cells were maintained in a 37 °C, 5 % CO₂ incubator for 4 h then change to EGM2- MV. Cells were harvested for qPCR and Western blot 48 h after transfection.

#### Analysis of 10x Multiome RNA+ATAC profiling of BPD and Control Lung

RNA and ATAC reads from 10x single nucleus multiome RNA and ATAC sequencing were preprocessed using the Cell Ranger ARC software package v 2.0.0 (10x Genomics) and human reference hg38 (10x Genomics refdata-cellranger-arc- GRCh38-2020-A-2.0.0). Nuclei passed Cell Ranger ARC filters were used as input to Signac (ref) for the following RNA and ATAC quality controls, including (i) RNA data has > 500 genes, < 15,000 UMIs, and < 15 % mitochondrial reads, and (ii) ATAC data have 500-25,000 UMIs, < 2 nucleosome signal values, and > 3 transcription start site enrichment score. In addition, Scrublet was applied to the RNA data to identify and remove potential doublets.^55^ In total, 24,749 cells were included in the downstream analysis. SoupX was applied to each multiome sample to remove technical ambient RNA counts from the RNA data.^56^ ATAC peaks from individual samples were merged to a common peak set (n = 17,384 peaks) for accessibility counting using Signac. RNA and ATAC data from different samples were integrated using the Seurat’s weighted nearest neighbor (WNN) analysis framework. Specifically, we first performed data integration using RNA and ATAC profiling separately using reciprocal principal component analysis (RPCA) and reciprocal latent semantic indexing (RLSI) analysis, respectively, and then we performed the integration using both the RNA and ATAC information by combining RPCA and RLSI dimensions using the WNN analysis. The WNN dimensions were used for UMAP projection and clustering analysis. Cell clusters were identified using the Leiden algorithm with default resolution. Cell clusters were annotated to cell types based were selective expressions of known cell type marker genes. Differential gene expression between two cell groups was performed using Seurat FindMarkers with two-tailed Wilcoxon rank sum test. Robust gene differential expression in BPD vs.

control gCap cells was identified using the following approach: (1) first, we compared gene expression in gCap cells between each BPD and control sample pair using Seurat FindMarkers function using two-tailed Wilcoxon rank sum test. (2) We also tested gene differential expression between all BPD vs. control gCap cells. In each comparison in (1) and (2), differential expression with the following criteria was considered significant: fold change >=1.5, expression percentage >=20%, and p value <0.05. (3) Genes significantly differentially expressed in (2) and in at least 4 out of 6 comparisons in (1) were included in the differential expression gene list. Ribosomal and mitochondrial genes were excluded.

#### Analysis of 10x Visium Spatial Transcriptomics Profiling of BPD and Control Lung

10x Visium spatial transcriptomics (ST) data were processed and quantified using 10x Space Ranger (v1.3.0) with hg38 reference genome (10x Genomics refdata-gex-GRCh38-2020-A). ST spots passed the Space Ranger filters were used as input to Seurat (v4.2.0) for downstream quality controls based on number of expressed genes and percentage of UMIs mapped to mitochondrial genes.^57^ Gene expression normalization was performed using Seurat SCTransform analysis. Cell type deconvolution was performed using Cell2location^58^ using both LungMAP human lung CellRef and the present multiomic data as single cell references.

We integrated single cell and spatial transcriptomics data to identify ligand-receptor pairs to/from gCap-C2 cells in BPD. First, we performed CellChat analysis using the single cell multiome profiling of each BPD sample and identified significant ligand-receptor (L-R) interactions associated with gCap-C2 cells in at least two of the three BPD samples. These candidate L-R pairs were subsequently subjected to spatial colocalization analysis using the 10x ST data of BPD samples. Each ST sample, we integrated the cell2location-deconvolution results using both the LungMAP CellRef and the present multiome data and identified *NTRK2*^+^ gCap C2 spots as those satisfying the following criteria: *NTRK2* expression >0, the estimated abundance of gCap-C2 >0.5 in the cell2location deconvolution using both references, and the proportion of gCap-C2 abundance >0.05 in the cell2location deconvolution using both references. Two BPD samples with >5 *NTRK2*^+^ gCap C2 spots were included in the downstream analysis. We then determined the 1-hop neighbors for each *NTRK2*^+^ gCap-C2 spot in the selected samples. For each CellChat identified LR pair from gCap-C2, we considered its ligand and receptor co-localized in a NTRK2+ gCap-C2 spot if the ligand is expressed in the spot and the receptor is expressed in the spot or its 1-hop neighbors. For each CellChat identified LR pair to gCap-C2, we considered its ligand and receptor co-localized in a NTRK2+ gCap-C2 spot if the receptor is expressed in the spot and the ligand is expressed in the spot or its 1-hop neighbors. LR pairs co-localized in at least 10% of the NTRK2+ gCap-C2 spots were selected. LR pairs from gCap-C2 were prioritized if the expression of the ligands was increased (fold change >=1.2, p value <0.05, and percentage>20%) in BPD vs. control gCap cells in the multiple profiling.

#### RNA scope staining

RNA in situ hybridization was performed for target detection on fresh 4 % PFA-fixed paraffin embedded 5□μm tissue sections using RNAscope Multiplex Fluorescent Reagent Kit Version 2 (Advanced Cell Diagnostics, Newark, CA, USA) according to the manual. Firstly, tissue sections were baked for 1□h at 60□°C, then deparaffinized and treated with hydrogen peroxide for 10□min at room temperature (RT). Target retrieval was performed for 15□min at 98□°C, followed by protease plus treatment for 15□min at 40□°C. All probes were hybridized for 2□h at 40□°C followed by signal amplification and development of HRP channels was done according to manual. The RNA-scope probes were used in the study: MM-*Ntrk2*-C2, Hs-*NTRK2*-O3-C2, Hs- *NTRK2*, Hs-*NTRK2*-O1-C1, Hs-*NTRK2*-O3-C1, MM-*Bdnf*-C3, HS-*PECAM1-*O1-C3, Hs-*AGER* - transcript variant 1, Hs-*AGER*-C3. Sections were counterstained with DAPI and mounted with ProLong Gold Antifade Mountant (Invitrogen).

#### Immunofluorescence (IF) Staining

For FFPE embedded sample, the sections (5 µm) from organoids were deparaffinized and rehydrated with a series of alcohol solutions of decreasing concentrations (5 min in each solution, 100 %, 95 %, 70 %, and 50 %) and kept in PBS for 30 min and moved to antigen retrieval process. For OCT embedded sample, the section (7 µm) from organoids were washed in PBS for 5 min and then fixed in 10 % Neutral buffered formalin (NBF) for 10 min. Slides were washed thrice for 5 min in PBS and then moved to antigen retrieval. Antigen retrieval was performed on sections immersed in citrate 10 mM in PBS, pH 6 and microwave heated (7 min once at 650 W and 5 min twice at 350 W). Sections were washed for 10 min in PBS and then incubated for 60 min at room temperature (RT) with blocking buffer (PBS containing 4 % donkey serum and 0.1 % Triton X-100). Then the sections were incubated with the primary antibodies diluted in blocking buffer at 4 °C overnight.

After three washes (5 min) in PBS, slides were incubated for 60 min at RT with secondary antibody diluted in blocking buffer. Slides were then washed three times for 5 min in PBS and mounted in aqueous mounting medium with DAPI. The first antibodies used are listed in Key Resource Table.

#### Imaging

Brightfield H&E images were acquired on a Nikon Eclipse NiE Upright Widefield Microscope (Nikon DS-Fi3 Camera—with a Plan Apo VC 20 × DIC N2 objective). Fluorescent images were acquired on Nikon A1 inverted LUNV and Nikon A1R inverted LUNV confocal microscopes using the following objectives: Plan Apo λ 10 ×, Plan Apo λ 20 ×, Apo LWD 20 × WI λS (water immersion), Apo LWD 40x WI λS DIC N2 (water immersion), and SR HP Plan Apo λ S 100 ×C Sil (silicone immersion). Second harmonic generation was performed to visualize fibrillar collagens I and II using a Nikon FN1 Upright Multiphoton microscope using the following objectives: Plan Apo VC 20 × DIC N2 and Apo LWD 25 × 1.10W DIC N2. Images were processed in Nikon Elements with minimal global adjustment of LUTs for acquired channels.

#### IVIS Imaging

Newborn mice were intravenously injected with Luciferase mRNA-loaded LNPs. The control group received empty LNPs. After 24 h, the mice were administered D-Luciferin (150 mg/kg, intraperitoneally), and tissues were collected post-euthanasia. Bioluminescence imaging was performed using an IVIS Lumina system.

#### Quantitative reverse-transcription PCR

Our detailed protocol was previously published.^59^ Briefly, total RNA was extracted, purified, and quantified for reverse transcription using High Capacity RNA to cDNA Kit (Applied Biosystems) according to the manufacturer’s instructions. qPCR was carried out using 5 ng cDNA and 6 μL SYBR green master mix (Applied Biosystems). Primers were listed in Table 3. Each measurement was performed in triplicate.

#### Western Blot

HPMVEC cells were subjected to mRNA transfection as described, followed by lysate preparation. Subsequent experimental procedures were performed according to standard protocols . Primary and secondary antibodies employed for Western blot analysis are listed in the Materials section. Protein bands were visualized using SuperSignal™ West Pico PLUS Chemiluminescent Substrate (Thermo Fisher

Scientific) and imaged using two independent systems: ChemiDoc™ Imaging System (Bio-Rad Laboratories, Hercules, CA, USA). Protein expression levels were quantified through densitometric analysis using ImageJ software (Version 1.53k; National Institutes of Health, Bethesda, MD, USA) with normalization to GAPDH as the loading control.

#### Rho GTPase activity assay

RhoA, Rac1, and Cdc42 activity were assessed in lysates from human pulmonary microvascular endothelial cells (HPMVECs) transfected with *GFP*, *NTRK2-FL*, or *NTRK2-T1*mRNA. Following transfection, cells were serum-starved for 24 h and subsequently stimulated with or without 25 ng/mL BDNF for 30 min to evaluate the effects on RhoA, Rac1, and Cdc42 activation.

After stimulation, cells were lysed in ice-cold lysis buffer provided in the G-LISA™ kits (Cytoskeleton, USA). Lysates were clarified by centrifugation at 10,000 rpm for 1 min at 4 °C, and protein concentrations were quantified using the BCA assay. To minimize GTP hydrolysis, total cell lysates were immediately flash-frozen in liquid nitrogen. Protein concentrations were standardized to 2 mg/mL for subsequent analysis.

RhoA, Rac1, and Cdc42 activities were measured using the G-LISA™ RhoA/Rac1/Cdc42 GTPase Activation Assay Bundle following the manufacturer’s instructions. Briefly, lysates were added to pre-coated wells specific to active RhoA, Rac1, and Cdc42. Primary antibodies targeting each GTPase were applied, followed by HRP-conjugated secondary antibodies. Signal detection was performed as per the kit protocol, and activity levels were normalized to the GFPgroup, expressed as fold changes relative to unstimulated controls.

#### Intracellular Ca^2+^ detection

Intracellular calcium levels were assessed using the Fluo-4 AM calcium indicator (Thermo Fisher Scientific) in HPMVECs. Cells seeded in 35 mm confocal dishes were incubated with 5 μM Fluo-4 AM in EGM-2 medium at 37 °C for 30 min in the dark. Following incubation, excess dye was removed by washing with Hank’s Balanced Salt Solution (HBSS), and cells were allowed to undergo a 10 min de-esterification period at room temperature to ensure complete intracellular conversion of Fluo-4 AM to its active form.

For live-cell imaging, the medium was replaced with Ca²⁺-free HBSS to eliminate extracellular calcium influence. Calcium distribution within cells was visualized using a confocal microscope (Leica Stellaris), and mean fluorescence intensity (MFI) was quantified and compared across experimental groups.

#### BrdU assay

The proliferation assay was carried out using BrdU Cell Proliferation Kit and following the manufacturer’s instructions (Millipore Sigma) at 72 h after transfection.

#### Tube formation assay

HPMVEC were transfected with *NTRK2* mRNA starved in medium containing 0.2 % FBS overnight and seeded at the density of 20,000 cells per 24-well plate in growth factor reduced Matrigel (Corning). The formation of tubes was observed by phase-contrast microscopy (Nikon, Japan) and quantified by ImageJ in three randomly selected fields at 12 h afterward.

#### Trans-endothelial electrical resistance (TEER) and paracellular permeability assays

Endothelial barrier integrity was assessed by TEER and paracellular permeability in HPMVECs. Cells transfected with vector and *NTRK2-T1* mRNA for 48 h and then digested, were seeded onto collagen I--coated Transwell inserts (0.4 μm pore size, Corning, USA) at a density of 1 × 105 cells per insert and cultured for 6 h until reaching confluence. TEER was measured using an EVOM3 volt-ohmmeter (World Precision Instruments, USA) with STX-2-plus chopstick electrodes. Readings were taken at baseline and after experimental treatments, followed by stimulation with or without 25 ng/mL BDNF for 24 h and 48 h. TEER values (Ω·cm²) were calculated by subtracting background resistance from cell-free inserts and normalizing to the surface area of the Transwell membrane.

Paracellular permeability was assessed using FITC-dextran (4 kDa, Sigma-Aldrich, USA) and rhodamine B-dextran (70 kDa, Sigma-Aldrich, USA). After experimental treatments, 600 μL of 1 mg/mL dextran in HBSS was added to the lower chamber, and 100 μL of HBSS was collected from the upper and lower chamber after 45 min. Fluorescence intensity was measured using a microplate reader, and dextran flux was expressed as fold change relative to the control group.

#### RNA immunoprecipitation (RIP) method and analysis

RIP experiment was performed to assess the interaction between RBFOX2 protein and *NTRK2*mRNA in HPMVECs using the Magna RIP™ RNA-Binding Protein Immunoprecipitation Kit (Millipore, USA) according to the manufacturer’s protocol. Briefly, HPMVECs were lysed in Complete RIP Lysis Buffer (RIP Lysis Buffer supplemented with protease inhibitor and RNase-free water) to obtain cleared lysates. Protein A/G magnetic beads were pre-washed with RIP Wash Buffer and conjugated with either RBFOX2-specific antibody (Cat. A300-864A, Thermo Fisher, USA) or normal IgG (negative control) for 30 min at room temperature. The antibody-bound beads were then incubated with the cleared lysates overnight at 4°C with gentle rotation. After incubation, the beads were washed six times with RIP Wash Buffer to eliminate non-specific interactions. Bound RNA was eluted using Elution Buffer and treated with Proteinase K to degrade proteins. RNA was then purified using phenol-chloroform-isoamyl alcohol extraction, followed by ethanol precipitation. The enriched RNA was reverse transcribed into cDNA using High-Capacity cDNA Reverse Transcription Kit (Cat. 4368814, Thermo Fisher, USA). qPCR was performed using primers specific to *NTRK2-T1* pre-mRNA. Fold enrichment was calculated relative to the IgG control using the 2^^(-ΔΔCt)^ method. This approach enabled the assessment of the specific interaction between RBFOX2 and *NTRK2T1* pre-mRNA in HPMVECs.

#### IVIS Imaging

Newborn mice were intravenously injected with Luciferase mRNA-loaded LNPs. The control group received empty LNPs. After 24 h, the mice were administered D-Luciferin (150 mg/kg, intraperitoneally), and tissues were collected post-euthanasia. Bioluminescence imaging was performed using an IVIS Lumina system.

#### Cre mRNA-LNP distribution in lungs using Ai14 mice model (Flow cytometry)

The newborn Ai14 mice was transorbital injected with 1 ug/g Cre mRNA-LNP at P2 and P4. 10 mU/g of Heparin sodium (LEO Pharma INc., Thornhill, ON, Canada) was administered intraperitoneally for flow cytometry procedures. Lung was harvested at P7 and prepared into single cell suspension as previously described. BD Horizon™ Fixable Viability Stain 510 was used to gate live cells. APC anti-mouse CD31 Antibody (BioLegend), PE/Cyanine7 anti-mouse CD45 Antibody (Biolegend), Alexa Fluor488-CD324(eBioscience, Invitrogen) and Brilliant Violet 421-CD140a (biolegend)was used to mark different cell type. Flow cytometry was using BD LSR Fortessa.

### QUANTIFICATION AND STATISTICAL ANALYSIS

For all image quantification, ImageJ and Imaris software were used. The frequency distribution for each imaging object was plotted in GraphPad Prism 8.0 and the Gaussian curve was created by nonlinear regression of the frequency distribution. The vascular network parameters were calculated with Imaris software with the ‘Neurofilament’ module.

Normal distribution from each group was confirmed using c2 test before any comparison between groups. All data represent mean ± SEM unless otherwise specified. Statistical significance was determined by parametric tests (all data possesses equal variance *P* > 0.05): unpaired 2-tailed t-test, ANOVA (>2 groups) with Bonferroni/Šídák’s test as indicated; or non-parametric test: Mann-Whitney (2 groups). A p value of < 0.05 was considered significant. \**P* < 0.05, \*\**P* < 0.01, \*\*\**P* < 0.001. Statistical details can be found in the figure legends. n represents different specimens from different cell lines, biological repeats, or patients or control subjects. The exact information for the n was specified in the figure legends. All violin plot data were tested by Bonferroni correction test and FDR < 0.05 was regarded as significant. Analyses were carried out using GraphPad Prism 8.0. As for all the experiments, at least three independent experiments were performed to reach the minimal requirement for statistical significance, unless otherwise specified. Blinding or randomization was not performed unless otherwise specified. Exclusion is not applied in this study.

### ADDITIONAL RESOURCES

N/A

## REFERENCES

1. Northway, W.H., Rosan, R.C., and Porter, D.Y. (1967). Pulmonary disease following respirator therapy of hyaline-membrane disease. Bronchopulmonary dysplasia. N Engl J Med 276, 357–368.

2. Jobe, A.H., and Bancalari, E. (2001). Bronchopulmonary dysplasia. Am J Respir Crit Care Med 163, 1723–1729.

3. Stoll, B.J., Hansen, N.I., Bell, E.F., Shankaran, S., Laptook, A.R., Walsh, M.C., Hale, E.C., Newman, N.S., Schibler, K., Carlo, W.A., et al. (2010). Neonatal outcomes of extremely preterm infants from the NICHD Neonatal Research Network. Pediatrics 126, 443–456. 10.1542/peds.2009-2959.

4. Bancalari, E., and Claure, N. (2022). Importance and Challenges Associated with Oxygen Control in Premature Infants. J Pediatr 247, 8–9. 10.1016/j.jpeds.2022.05.042.

5. Check, J., Jensen, E.T., Skelton, J.A., Ambrosius, W.T., and O’Shea, T.M. (2020). Early growth outcomes in very low birth weight infants with bronchopulmonary dysplasia or fetal growth restriction. Pediatr Res 88, 601–604. 10.1038/s41390-020-0808-7.

6. Thébaud, B., Goss, K.N., Laughon, M., Whitsett, J.A., Abman, S.H., Steinhorn, R.H., Aschner, J.L., Davis, P.G., McGrath-Morrow, S.A., Soll, R.F., and Jobe, A.H. (2019). Bronchopulmonary dysplasia. Nat Rev Dis Primers 5, 78. 10.1038/s41572-019-0127-7.

7. Lim, G., Lee, B.S., Choi, Y.-S., Park, H.W., Chung, M.L., Choi, H.J., Kim, E.A.-R., and Kim, K.-S. (2015). Delayed Dexamethasone Therapy and Neurodevelopmental Outcomes in Preterm Infants with Bronchopulmonary Dysplasia. Pediatr Neonatol 56, 261–267. 10.1016/j.pedneo.2014.11.006.

8. Qin, G., Lo, J.W., Marlow, N., Calvert, S.A., Greenough, A., and Peacock, J.L. (2017). Postnatal dexamethasone, respiratory and neurodevelopmental outcomes at two years in babies born extremely preterm. PLoS One 12, e0181176. 10.1371/journal.pone.0181176.

9. Abman, S.H., Collaco, J.M., Shepherd, E.G., Keszler, M., Cuevas-Guaman, M., Welty, S.E., Truog, W.E., McGrath-Morrow, S.A., Moore, P.E., Rhein, L.M., et al. (2017). Interdisciplinary Care of Children with Severe Bronchopulmonary Dysplasia. J Pediatr 181. 10.1016/j.jpeds.2016.10.082.

10. Baker, C.D., Ryan, S.L., Ingram, D.A., Seedorf, G.J., Abman, S.H., and Balasubramaniam, V. (2009). Endothelial colony-forming cells from preterm infants are increased and more susceptible to hyperoxia. Am J Respir Crit Care Med 180, 454–461. 10.1164/rccm.200901-0115OC.

11. Cristea, A.I., Ren, C.L., Amin, R., Eldredge, L.C., Levin, J.C., Majmudar, P.P., May, A.E., Rose, R.S., Tracy, M.C., Watters, K.F., et al. (2021). Outpatient Respiratory Management of Infants, Children, and Adolescents with Post- Prematurity Respiratory Disease: An Official American Thoracic Society Clinical Practice Guideline. Am J Respir Crit Care Med 204, e115–e133. 10.1164/rccm.202110-2269ST.

12. Krishnan, U., Feinstein, J.A., Adatia, I., Austin, E.D., Mullen, M.P., Hopper, R.K., Hanna, B., Romer, L., Keller, R.L., Fineman, J., et al. (2017). Evaluation and Management of Pulmonary Hypertension in Children with Bronchopulmonary Dysplasia. J Pediatr 188. 10.1016/j.jpeds.2017.05.029.

13. Hansmann, G., Sallmon, H., Roehr, C.C., Kourembanas, S., Austin, E.D., and Koestenberger, M. (2021). Pulmonary hypertension in bronchopulmonary dysplasia. Pediatr Res 89, 446–455. 10.1038/s41390-020-0993-4.

14. 14. Jakkula, M., Le Cras, T.D., Gebb, S., Hirth, K.P., Tuder, R.M., Voelkel, N.F., and Abman, S.H. (2000). Inhibition of angiogenesis decreases alveolarization in the developing rat lung. Am J Physiol Lung Cell Mol Physiol 279, L600–L607.

15. Ladha, F., Bonnet, S., Eaton, F., Hashimoto, K., Korbutt, G., and Thébaud, B. (2005). Sildenafil improves alveolar growth and pulmonary hypertension in hyperoxia-induced lung injury. Am J Respir Crit Care Med 172, 750–756.

16. Thébaud, B., Ladha, F., Michelakis, E.D., Sawicka, M., Thurston, G., Eaton, F., Hashimoto, K., Harry, G., Haromy, A., Korbutt, G., and Archer, S.L. (2005). Vascular endothelial growth factor gene therapy increases survival, promotes lung angiogenesis, and prevents alveolar damage in hyperoxia- induced lung injury: evidence that angiogenesis participates in alveolarization. Circulation 112, 2477–2486.

17. Cantu, A., Cantu Gutierrez, M., Zhang, Y., Dong, X., and Lingappan, K. (2023). Endothelial to mesenchymal transition in neonatal hyperoxic lung injury: role of sex as a biological variable. Physiol Genomics 55, 345–354. 10.1152/physiolgenomics.00037.2023.

18. Ellis, L.V., Bywaters, J.D., and Chen, J. (2024). Endothelial deletion of p53 generates transitional endothelial cells and improves lung development during neonatal hyperoxia. bioRxiv.10.1101/2024.05.07.593014.

19. Zanini, F., Che, X., Knutsen, C., Liu, M., Suresh, N.E., Domingo-Gonzalez, R., Dou, S.H., Zhang, D., Pryhuber, G.S., Jones, R.C., et al. (2023). Developmental diversity and unique sensitivity to injury of lung endothelial subtypes during postnatal growth. iScience 26, 106097. 10.1016/j.isci.2023.106097.

20. Niethamer, T.K., Planer, J.D., Morley, M.P., Babu, A., Zhao, G., Basil, M.C., Cantu, E., Frank, D.B., Diamond, J.M., Nottingham, A.N., et al. (2025). Longitudinal single-cell profiles of lung regeneration after viral infection reveal persistent injury-associated cell states. Cell Stem Cell 32, 302–321.e306. 10.1016/j.stem.2024.12.002.

21. Raslan, A.A., Pham, T.X., Lee, J., Kontodimas, K., Tilston-Lunel, A., Schmottlach, J., Hong, J., Dinc, T., Bujor, A.M., Caporarello, N., et al. (2024). Lung injury-induced activated endothelial cell states persist in aging- associated progressive fibrosis. Nat Commun 15, 5449. 10.1038/s41467-024- 49545-x.

22. Travaglini, K.J., Nabhan, A.N., Penland, L., Sinha, R., Gillich, A., Sit, R.V., Chang, S., Conley, S.D., Mori, Y., Seita, J., et al. (2020). A molecular cell atlas of the human lung from single-cell RNA sequencing. Nature 587, 619–625. 10.1038/s41586-020-2922-4.

23. Middlemas, D.S., Lindberg, R.A., and Hunter, T. (1991). trkB, a neural receptor protein-tyrosine kinase: evidence for a full-length and two truncated receptors. Mol Cell Biol 11, 143–153. 10.1128/mcb.11.1.143-153.1991.

24. Klein, R., Nanduri, V., Jing, S.A., Lamballe, F., Tapley, P., Bryant, S., Cordon- Cardo, C., Jones, K.R., Reichardt, L.F., and Barbacid, M. (1991). The trkB tyrosine protein kinase is a receptor for brain-derived neurotrophic factor and neurotrophin-3. Cell 66, 395–403. 10.1016/0092-8674(91)90628-c.

25. Xia, Y., Wang, Z.-H., Liu, P., Edgington-Mitchell, L., Liu, X., Wang, X.-C., and Ye, K. (2021). TrkB receptor cleavage by delta-secretase abolishes its phosphorylation of APP, aggravating Alzheimer’s disease pathologies. Mol Psychiatry 26, 2943–2963. 10.1038/s41380-020-00863-8.

26. Chaudhuri, D., Sasaki, K., Karkar, A., Sharif, S., Lewis, K., Mammen, M.J., Alexander, P., Ye, Z., Lozano, L.E.C., Munch, M.W., et al. (2021). Corticosteroids in COVID-19 and non-COVID-19 ARDS: a systematic review and meta-analysis. Intensive Care Med 47, 521–537. 10.1007/s00134-021-06394-2.

27. Omar, S.A., Abdul-Hafez, A., Ibrahim, S., Pillai, N., Abdulmageed, M., Thiruvenkataramani, R.P., Mohamed, T., Madhukar, B.V., and Uhal, B.D. (2022). Stem-Cell Therapy for Bronchopulmonary Dysplasia (BPD) in Newborns. Cells 11. 10.3390/cells11081275.

28. Pattwell, S.S., Arora, S., Cimino, P.J., Ozawa, T., Szulzewsky, F., Hoellerbauer, P., Bonifert, T., Hoffstrom, B.G., Boiani, N.E., Bolouri, H., et al. (2020). A kinase-deficient NTRK2 splice variant predominates in glioma and amplifies several oncogenic signaling pathways. Nat Commun 11, 2977. 10.1038/s41467-020-16786-5.

29. Riccetti, M.R., Ushakumary, M.G., Waltamath, M., Green, J., Snowball, J., Dautel, S.E., Endale, M., Lami, B., Woods, J., Ahlfeld, S.K., and Perl, A.-K.T. (2022). Maladaptive functional changes in alveolar fibroblasts due to perinatal hyperoxia impair epithelial differentiation. JCI Insight 7. 10.1172/jci.insight.152404.

30. Tomassoni-Ardori, F., Fulgenzi, G., Becker, J., Barrick, C., Palko, M.E., Kuhn, S., Koparde, V., Cam, M., Yanpallewar, S., Oberdoerffer, S., and Tessarollo, L. (2019). Rbfox1 up-regulation impairs BDNF-dependent hippocampal LTP by dysregulating TrkB isoform expression levels. Elife 8. 10.7554/eLife.49673.

31. Madisen, L., Zwingman, T.A., Sunkin, S.M., Oh, S.W., Zariwala, H.A., Gu, H., Ng, L.L., Palmiter, R.D., Hawrylycz, M.J., Jones, A.R., et al. (2010). A robust and high-throughput Cre reporting and characterization system for the whole mouse brain. Nat Neurosci 13, 133–140. 10.1038/nn.2467.

32. Caporarello, N., Lee, J., Pham, T.X., Jones, D.L., Guan, J., Link, P.A., Meridew, J.A., Marden, G., Yamashita, T., Osborne, C.A., et al. (2022). Dysfunctional ERG signaling drives pulmonary vascular aging and persistent fibrosis. Nat Commun 13, 4170. 10.1038/s41467-022-31890-4.

33. Schupp, J.C., Adams, T.S., Cosme, C., Raredon, M.S.B., Yuan, Y., Omote, N., Poli, S., Chioccioli, M., Rose, K.-A., Manning, E.P., et al. (2021). Integrated Single-Cell Atlas of Endothelial Cells of the Human Lung. Circulation 144, 286–302. 10.1161/CIRCULATIONAHA.120.052318.

34. Guo, S., Kim, W.J., Lok, J., Lee, S.-R., Besancon, E., Luo, B.-H., Stins, M.F., Wang, X., Dedhar, S., and Lo, E.H. (2008). Neuroprotection via matrix-trophic coupling between cerebral endothelial cells and neurons. Proc Natl Acad Sci U S A 105, 7582–7587. 10.1073/pnas.0801105105.

35. Nie, X., Shen, C., Tan, J., Wu, Z., Wang, W., Chen, Y., Dai, Y., Yang, X., Ye, S., Chen, J., and Bian, J.-S. (2020). Periostin: A Potential Therapeutic Target For Pulmonary Hypertension? Circ Res 127, 1138–1152. 10.1161/CIRCRESAHA.120.316943.

36. Navaratna, D., Fan, X., Leung, W., Lok, J., Guo, S., Xing, C., Wang, X., and Lo, E.H. (2013). Cerebrovascular degradation of TRKB by MMP9 in the diabetic brain. J Clin Invest 123, 3373–3377. 10.1172/JCI65767.

37. Totoson, P., Pedard, M., Marie, C., and Demougeot, C. (2018). Activation of endothelial TrkB receptors induces relaxation of resistance arteries. Vascul Pharmacol 106, 46–53. 10.1016/j.vph.2018.02.005.

38. Dalton, J.E., Glover, A.C., Hoodless, L., Lim, E.-K., Beattie, L., Kirby, A., and Kaye, P.M. (2015). The neurotrophic receptor Ntrk2 directs lymphoid tissue neovascularization during Leishmania donovani infection. PLoS Pathog 11, e1004681. 10.1371/journal.ppat.1004681.

39. Fork, C., Gu, L., Hitzel, J., Josipovic, I., Hu, J., SzeKa Wong, M., Ponomareva, Y., Albert, M., Schmitz, S.U., Uchida, S., et al. (2015). Epigenetic Regulation of Angiogenesis by JARID1B-Induced Repression of HOXA5. Arteriosclerosis, Thrombosis, and Vascular Biology 35, 1645–1652. 10.1161/ATVBAHA.115.305561.

40. Liao, Y., Wang, C., Yang, Z., Liu, W., Yuan, Y., Li, K., Zhang, Y., Wang, Y., Shi, Y., Qiu, Y., et al. (2020). Dysregulated Sp1/miR-130b-3p/HOXA5 axis contributes to tumor angiogenesis and progression of hepatocellular carcinoma. Theranostics 10, 5209–5224. 10.7150/thno.43640.

41. Verma, S.K., Deshmukh, V., Thatcher, K., Belanger, K.K., Rhyner, A.M., Meng, S., Holcomb, R.J., Bressan, M., Martin, J.F., Cooke, J.P., et al. (2022). RBFOX2 is required for establishing RNA regulatory networks essential for heart development. Nucleic Acids Res 50, 2270–2286. 10.1093/nar/gkac055.

42. Wei, C., Qiu, J., Zhou, Y., Xue, Y., Hu, J., Ouyang, K., Banerjee, I., Zhang, C., Chen, B., Li, H., et al. (2015). Repression of the Central Splicing Regulator RBFox2 Is Functionally Linked to Pressure Overload-Induced Heart Failure. Cell Reports 10, 1521–1533. 10.1016/j.celrep.2015.02.013.

43. Zhou, Y., Fan, J., Zhu, H., Ji, L., Fan, W., Kapoor, I., Wang, Y., Wang, Y., Zhu, G., and Wang, J. (2017). Aberrant Splicing Induced by Dysregulated Rbfox2 Produces Enhanced Function of CaV1.2 Calcium Channel and Vascular Myogenic Tone in Hypertension. Hypertension 70, 1183–1192. 10.1161/HYPERTENSIONAHA.117.09301.

44. Li, L., Xie, S., and Deng, W. (2025). RNA binding proteins: Mechanistic considerations and perspectives in controlling cardiovascular diseases. European Journal of Pharmacology 987, 177101. 10.1016/j.ejphar.2024.177101.

45. Tomassoni-Ardori, F., Palko, M.E., Galloux, M., and Tessarollo, L. (2025). Clusters of deep intronic RbFox motifs embedded in large assembly of splicing regulators sequences regulate alternative splicing. bioRxiv. 10.1101/2024.08.19.608686.

46. Paris, A.J., Hayer, K.E., Oved, J.H., Avgousti, D.C., Toulmin, S.A., Zepp, J.A., Zacharias, W.J., Katzen, J.B., Basil, M.C., Kremp, M.M., et al. (2020). STAT3- BDNF-TrkB signalling promotes alveolar epithelial regeneration after lung injury. Nat Cell Biol 22, 1197–1210. 10.1038/s41556-020-0569-x.

47. Conte, G., Costabile, G., Baldassi, D., Rondelli, V., Bassi, R., Colombo, D., Linardos, G., Fiscarelli, E.V., Sorrentino, R., Miro, A., et al. (2022). Hybrid Lipid/Polymer Nanoparticles to Tackle the Cystic Fibrosis Mucus Barrier in siRNA Delivery to the Lungs: Does PEGylation Make the Difference? ACS Appl Mater Interfaces 14, 7565–7578. 10.1021/acsami.1c14975.

48. Somu Naidu, G., Rampado, R., Sharma, P., Ezra, A., Kundoor, G.R., Breier, D., and Peer, D. (2025). Ionizable Lipids with Optimized Linkers Enable Lung- Specific, Lipid Nanoparticle-Mediated mRNA Delivery for Treatment of Metastatic Lung Tumors. ACS Nano 19, 6571–6587.

49. 10.1021/acsnano.4c18636.

49. Zhang, L., More, K.R., Ojha, A., Jackson, C.B., Quinlan, B.D., Li, H., He, W., Farzan, M., Pardi, N., and Choe, H. (2023). Effect of mRNA-LNP components of two globally-marketed COVID-19 vaccines on efficacy and stability. npj Vaccines 8, 156. 10.1038/s41541-023-00751-6.

50. Rowe, R.G., and Daley, G.Q. (2019). Induced pluripotent stem cells in disease modelling and drug discovery. Nature Reviews Genetics 20, 377–388. 10.1038/s41576-019-0100-z.

51. Sayed, N., Wong, W.T., Ospino, F., Meng, S., Lee, J., Jha, A., Dexheimer, P., Aronow, B.J., and Cooke, J.P. (2015). Transdifferentiation of human fibroblasts to endothelial cells: role of innate immunity. Circulation 131, 300–309. 10.1161/circulationaha.113.007394.

52. Meng, S., Lv, J., Chanda, P.K., Owusu, I., Chen, K., and Cooke, J.P. (2020). Reservoir of Fibroblasts Promotes Recovery From Limb Ischemia. Circulation 142, 1647–1662. 10.1161/circulationaha.120.046872.

53. Radmand, A., Lokugamage, M.P., Kim, H., Dobrowolski, C., Zenhausern, R., Loughrey, D., Huayamares, S.G., Hatit, M.Z.C., Ni, H., Del Cid, A., et al. (2023). The Transcriptional Response to Lung-Targeting Lipid Nanoparticles in Vivo. Nano Lett 23. 10.1021/acs.nanolett.2c04479.

54. Wimmer, R.A., Leopoldi, A., Aichinger, M., Wick, N., Hantusch, B., Novatchkova, M., Taubenschmid, J., Hämmerle, M., Esk, C., Bagley, J.A., et al. (2019). Human blood vessel organoids as a model of diabetic vasculopathy. Nature 565, 505–510. 10.1038/s41586-018-0858-8.

55. Wolock, S.L., Lopez, R., and Klein, A.M. (2019). Scrublet: Computational Identification of Cell Doublets in Single-Cell Transcriptomic Data. Cell Syst 8. 10.1016/j.cels.2018.11.005.

56. Young, M.D., and Behjati, S. (2020). SoupX removes ambient RNA contamination from droplet-based single-cell RNA sequencing data. Gigascience 9. 10.1093/gigascience/giaa151.

57. Hao, Y., Hao, S., Andersen-Nissen, E., Mauck, W.M., Zheng, S., Butler, A., Lee, M.J., Wilk, A.J., Darby, C., Zager, M., et al. (2021). Integrated analysis of multimodal single-cell data. Cell 184. 10.1016/j.cell.2021.04.048.

58. Kleshchevnikov, V., Shmatko, A., Dann, E., Aivazidis, A., King, H.W., Li, T., Elmentaite, R., Lomakin, A., Kedlian, V., Gayoso, A., et al. (2022). Cell2location maps fine-grained cell types in spatial transcriptomics. Nat Biotechnol 40, 661–671. 10.1038/s41587-021-01139-4.

59. Gu, M., Shao, N.-Y., Sa, S., Li, D., Termglinchan, V., Ameen, M., Karakikes, I., Sosa, G., Grubert, F., Lee, J., et al. (2017). Patient-Specific iPSC-Derived Endothelial Cells Uncover Pathways that Protect against Pulmonary Hypertension in BMPR2 Mutation Carriers. Cell Stem Cell 20. 10.1016/j.stem.2016.08.019.

