## Supplemental images for "Isoform-Specific Roles of NTRK2 in Pulmonary Vascular Regeneration"

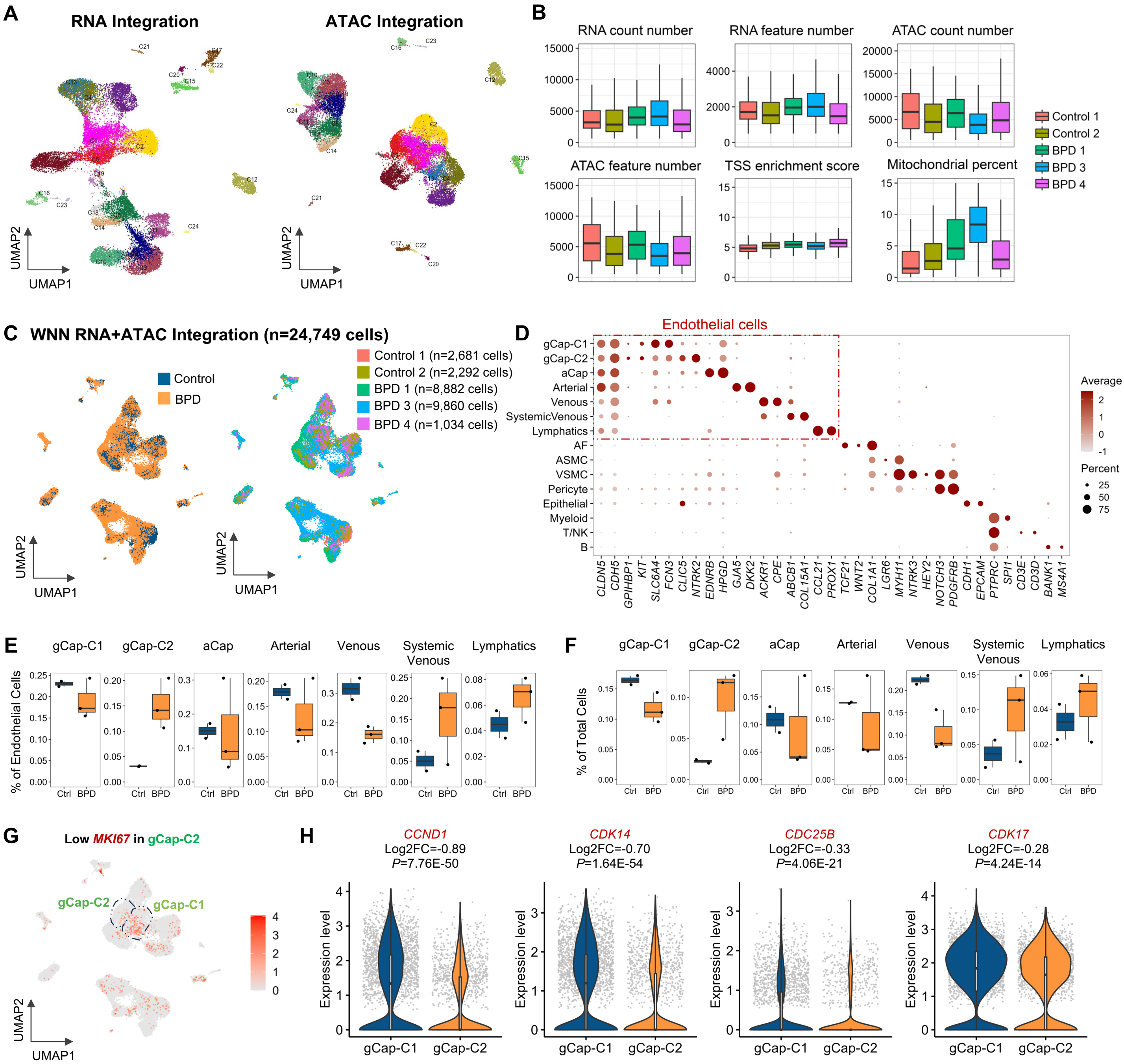


**Figure S1: Analysis of 10x multiome profiling of human BPD and control lung samples.**

**(A)** UMAP plots showing data integration using RNA (left) and ATAC (right) profiles independently. Cells in both the left and right panels were colored based on the same clustering result identified using the weighted nearest neighbor (WNN) analysis in (C), which integrated both RNA and ATAC patterns.

**(B)** Boxplots showing RNA UMI count, RNA feature count, ATAC UMI count, ATAC feature count, TSS enrichment score and mitochondria percentage in the cells of each multiome sample.

**(C)** UMAP plots showing data integration using both the RNA and ATAC patterns in (A) using the Seurat's weighted nearest neighbor (WNN) analysis method. Left: cells colored by condition. Right: cells colored by the sample.

**(D)** Dotplot showing the selective expression of marker genes in their corresponding cell types identified in the multiome analysis.

**(E)** Boxplots showing the percentage of each endothelial cell type among all endothelial cells in each multiome sample.

**(F)** Boxplots showing the percentage of each endothelial cell type among all cells in each multiome sample.

**(G)** UMAP plot showing expression of *MKI67* in all cells. Cells with positive *MKI67* expression (red) were plotted on top of cells with zero *MKI67* expression (grey).

**(H)** Violin and box plots showing expression of *CCND1*, *CDK14*, *CDC25B* and *CDK17* in gCap-C1 and gCap-C2 cells. P values and fold change (FC) values were obtained from tests of gene expression in gCap-C2 vs. gCap-C1 cells using Seurat FindMarkers function using two-tailed Wilcoxon rank sum test.

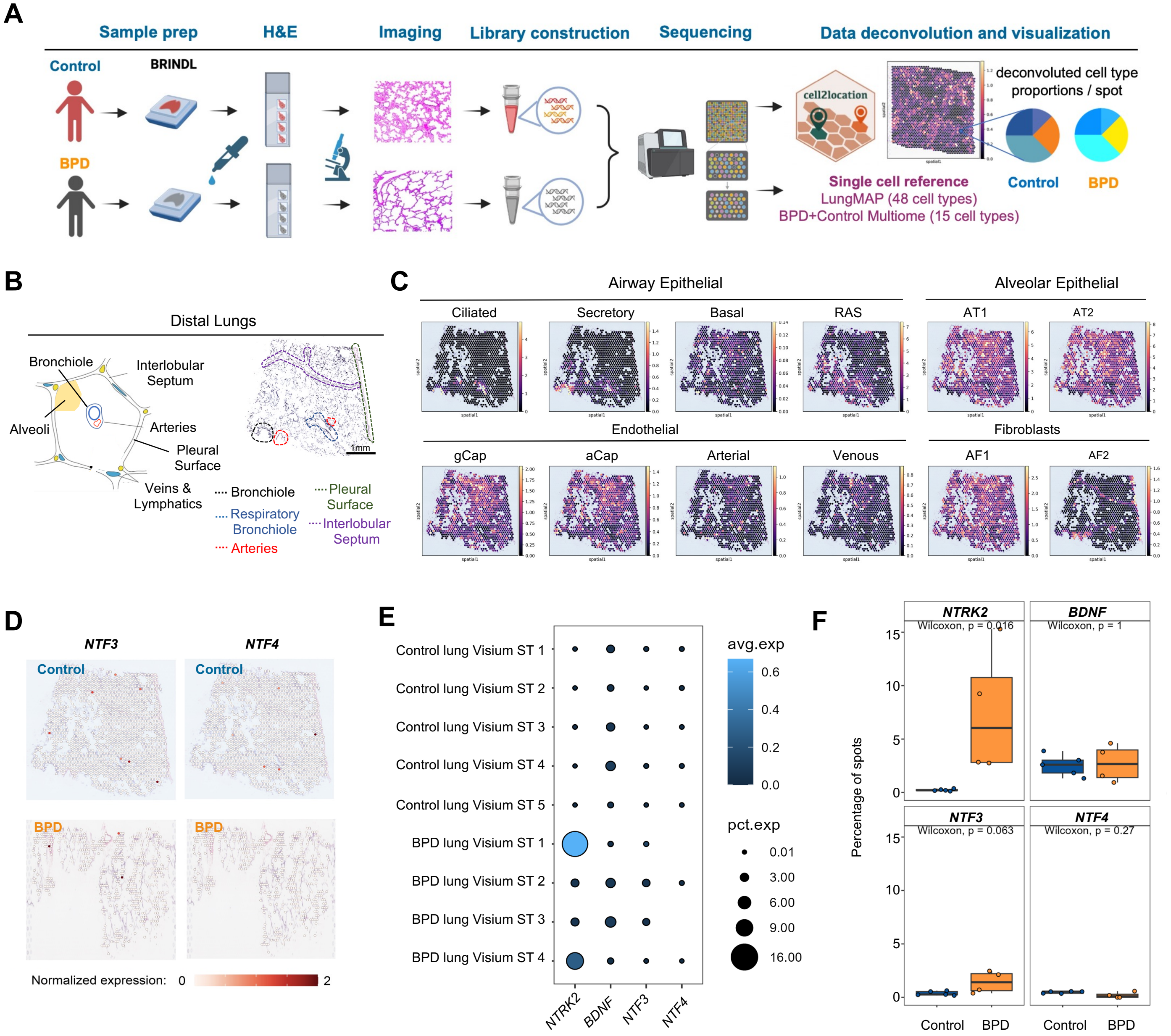


**Figure S2: Processing and analysis of 10x Visium spatial transcriptomics (ST) profiling of human BPD and control lung samples.**

(**A**) Schematic workflow of 10x Visium ST analysis of human BPD distal lung samples and age matched controls. Cell2location was used for the cell type deconvolution analysis using two single cell references: 10x Multiome profiling in Figure 1A and the LungMAP human lung CellRef (48 cell types, Guo et al., Nature Communication 2023).

(**B**) Illustration (left) and H&E staining with histological annotation (right) of a control human distal lung sample used in the Visium ST profiling.

(**C**) Visualization of Cell2location estimation of cell type abundance in each spatial spot in the sample shown in (B). Cell type identities were learned using the LungMAP human lung CellRef. AT1: alveolar type 1 cells; AT2: alveolar type 2 cells; EC: endothelial cells; gCap: general capillary; aCap: aerocyte; AF1: alveolar fibroblast type 1; AF2: alveolar fibroblast type 2; RAS: respiratory airway secretory cells.

(**D**) Expression of *NTF3* and *NTF4* in ST of example BPD and control Visium ST samples.

(**E**) Dotplot showing the expression frequencies and levels of *NTRK2* and its ligands in all BPD and control Visium ST samples.

(**F**) Box plot visualization of the expression frequencies of *NTRK2* and its ligands in Visium ST of BPD and control samples. n = 5 Visium ST samples for the control group, n = 4 Visium ST samples for the BPD group. P value represent significance of difference in expression frequencies of each gene between BPD and control Visium ST samples. Tests were performed using two-tailed Wilcoxon rank sum test.

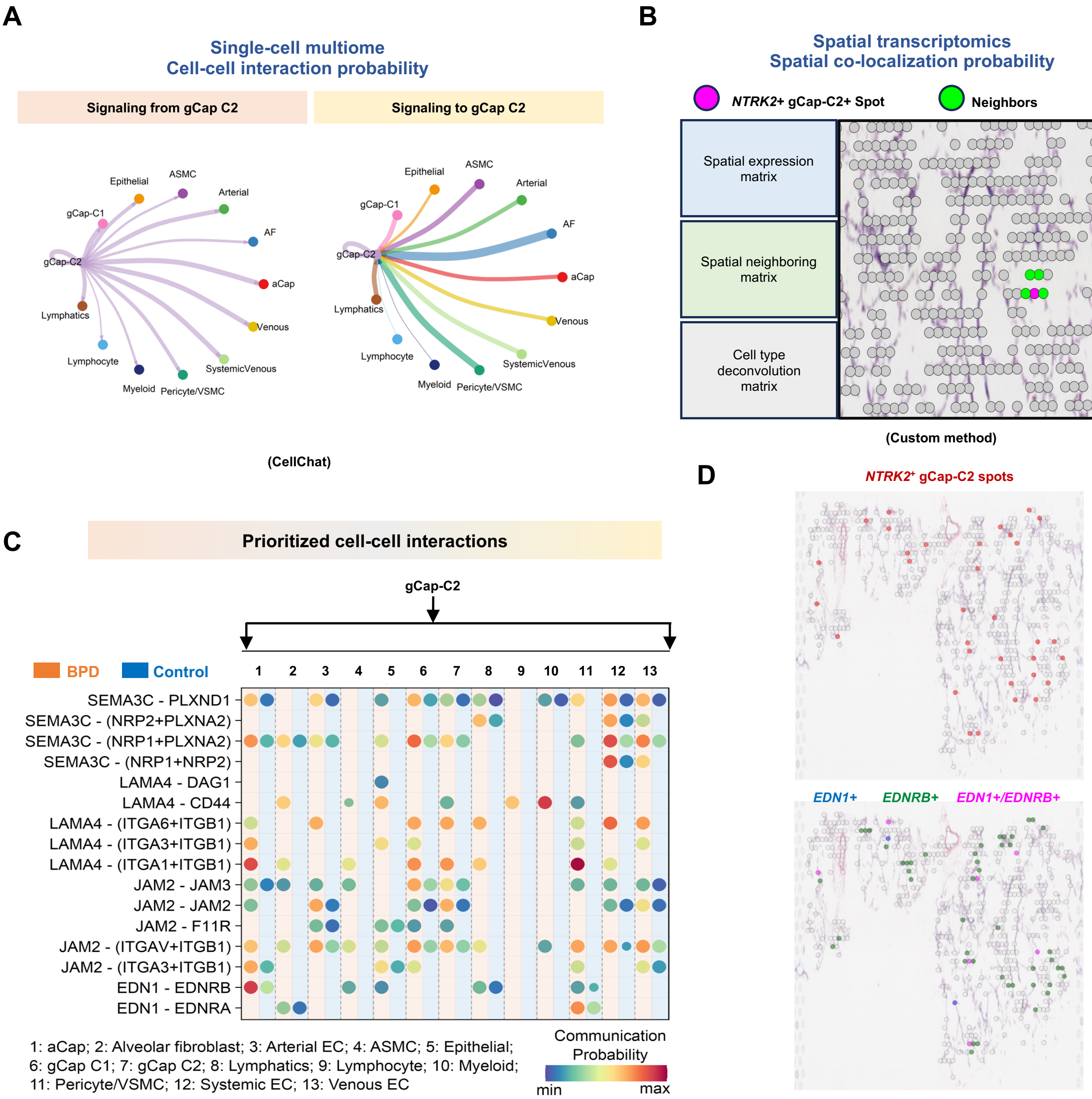


**Figure S3: Identification of gCap-C2 cell-cell interactions using an integrative analysis of single cell multiome and Visium ST data.**

(**A**) CellChat analysis was performed using single cell multiome data to identify ligand-receptor (LR) pairs to/from gCap-C2 cells in BPD samples.

(**B**) LR pairs identified in (A) were used as candidates for co-localization analysis using the Visium ST data. We integrated the information from spatial gene expression matrix, neighborhood matrix of spatial spots, and cell type deconvolution results to calculate a colocalization frequency for each LR pair near *NTRK2*+ gCap-C2 spots in ST of BPD samples, which measures the frequency of co-expression of a ligand-receptor pair within the same *NTRK2*+ gCap-C2 spot or in 1-hop neighbors (Methods).

(**C**) LR pairs from gCap-C2 cells to other cells shown in a CellChat analysis of BPD vs. control multiome data. The LR pairs were identified based on the analyses in (A) and (B). Shown are LR pairs with the expression of the ligands up-regulated (fold change >=1.2, expression percentage >=20%) in gCap cells in BPD vs. Control lung multiome data.

(**D**) Example of the co-localization (bottom) of *EDN1* and *EDNRB* near *NTRK2*+ gCap-C2 spots (top) in a BPD Visium ST sample.

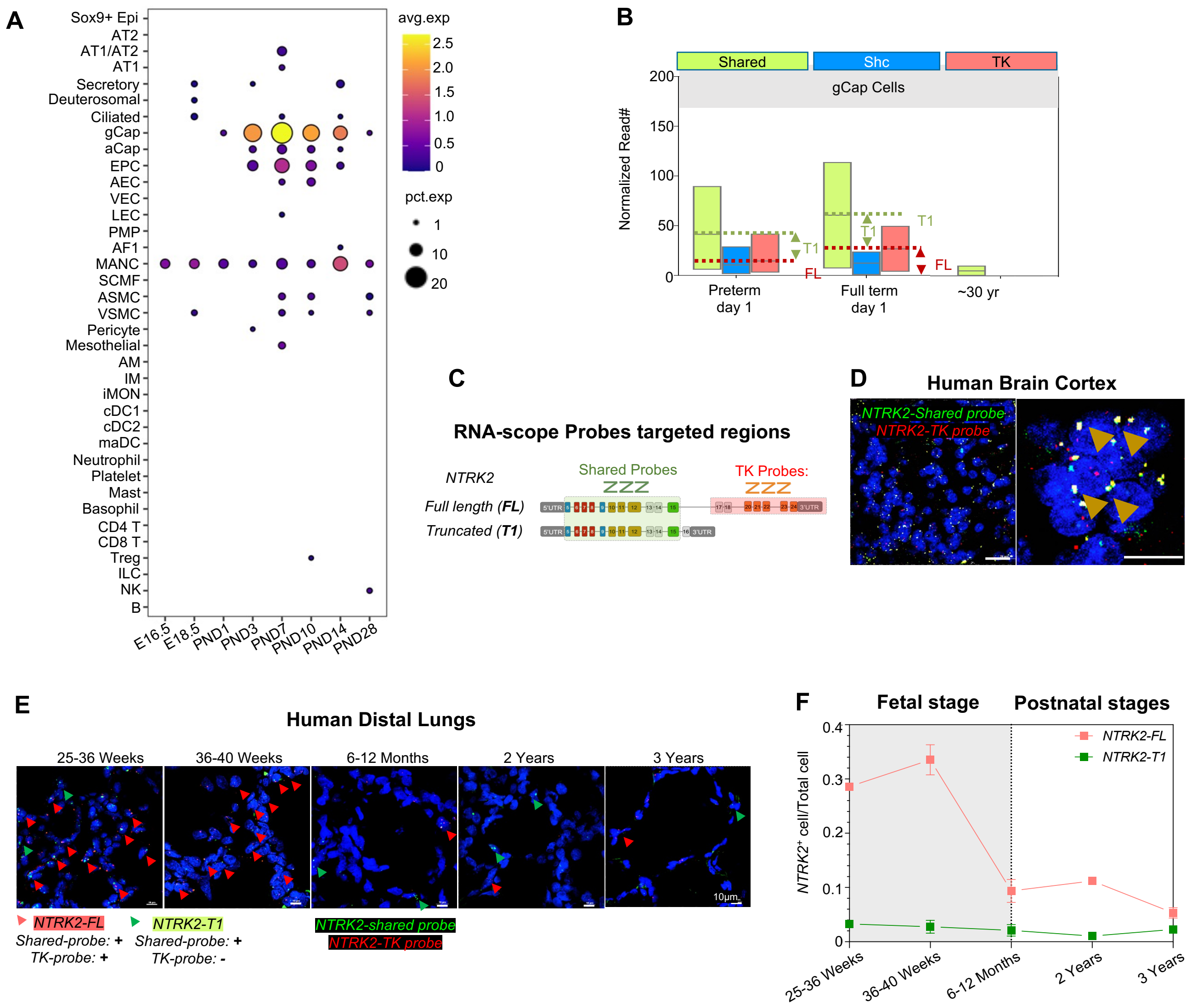


**Figure S4: *NTRK2* isoform expression patterns across multiple lung development stages.**

(**A**) Dotplot showing the expression frequencies and levels of *Ntrk2* in individual cells types in perinatal mouse lung development using single cell RNA-seq data in LungMAP mouse lung CellRef (Guo et al., Nature Communications 2023).

(**B**) Bar graph showing normalized scRNAseq reads of different region of *NTRK2* among gCap cells Preterm day1, full term day1 and adult lungs (preterm and adult lung data were from PMID: 33164753. Full term data was from LungMAP).

**(C)** Demonstration figure of RNA-scope probes designed specific to shared region and tyrosine kinase region of *NTRK2*.

**(D)** Representative RNA-scope staining result of human brain cortex using *NTRK2-shared* probe(green) and *NTRK2-TK* probe(Red). Brown arrow pointing the overlapped signal of *NTRK2-shared* probe and *NTRK2-TK* probe, indicating *NTRK2 FL* mRNA. Scale bar 20μm (left), 10μm (right).

**(E)** Representative RNA-scope staining result of different stages of human distal from mid term pregnancy to 3 years old. *NTRK2-shared* (green), *NTRK2-TK* (red). Green arrow indicating *NTRK2-T1* mRNA and red arrow indicating *NTRK2-FL* mRNA. Scale bar, 10 μm.

**(F)** The percentage of *NTRK2-FL* positive cells or *NTRK2-T1* positive cell among all cells at different stages of lung development. Data present by mean ± SEM. n = 3 patients.

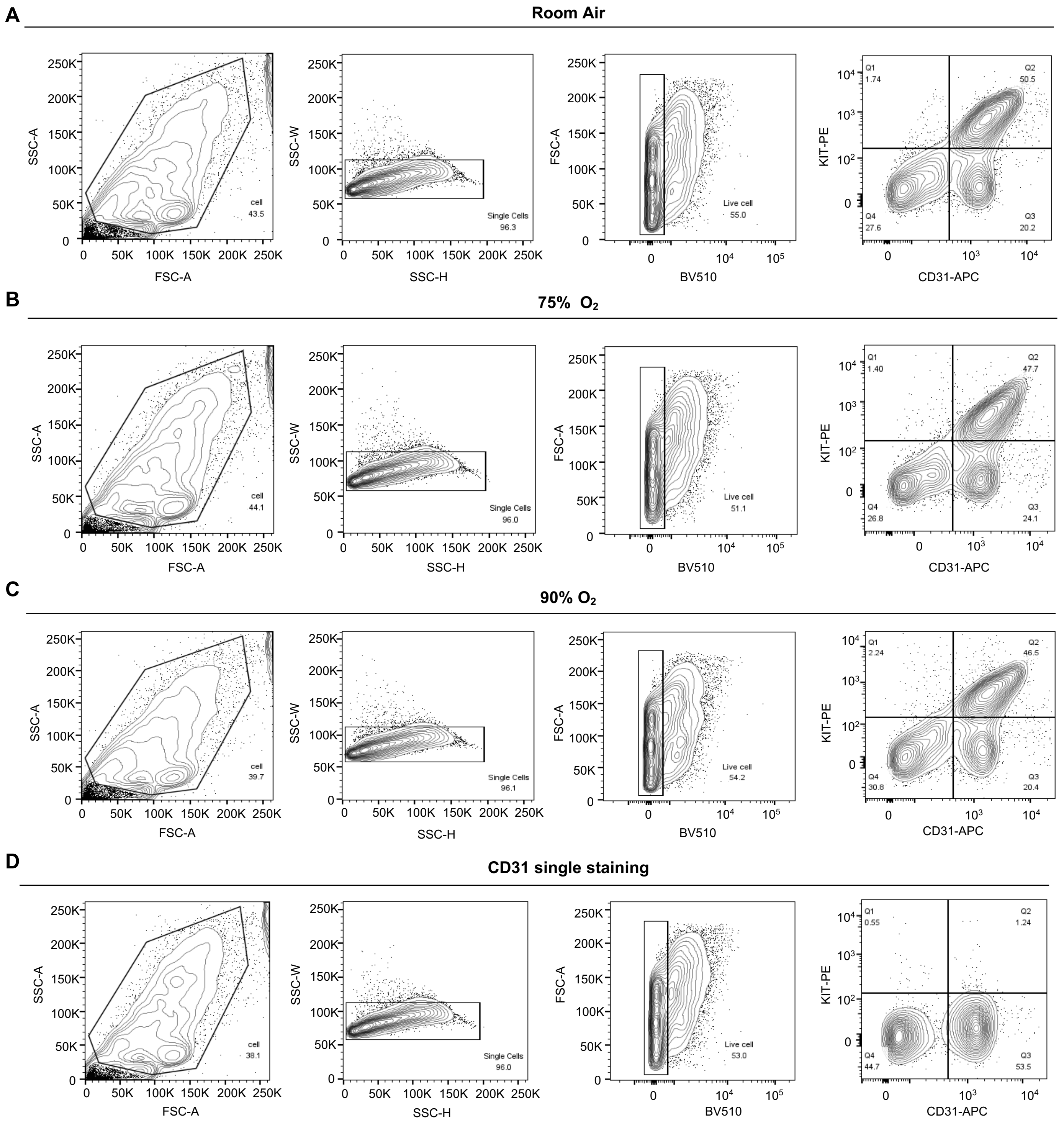


**Figure S5: Gating strategy of mouse gCap cells in distal lung at P10. (A-D)** Gating strategy of cell, singlets, live cell and gCap(Pecam-1/Kit+) in Room air (**A**), 75% $O_2$ (**B**), 90% $O_2$ (**C**), and CD31 single stain control (**D**).

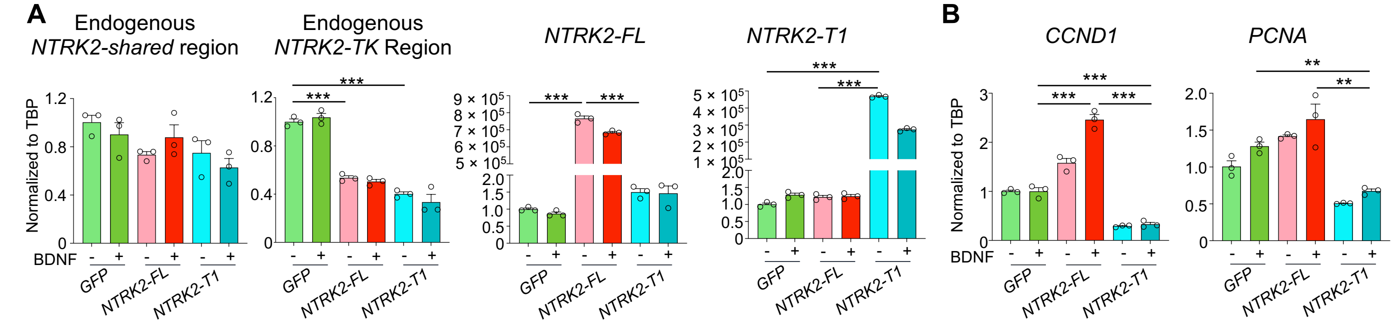


**Figure S6: Transfection of *NTRK2* isoforms in human pulmonary microvascular endothelial cells (HPMVECs).** (**A**) The expression of endogenous and exogenous *NTRK2-shared and NTRK2-TK* regions normalized to *GFP* group. Each dot refers to a biological repeat. Data were presented as mean ± SEM. ***$P < 0.001$ *NTRK2-FL* vs. *GFP*, *NTRK2-T1* vs. *GFP*, one-way ANOVA followed by Bonferroni's multiple comparisons (n = 3 repeats).
(**B**) The expression of *CCND1* and *PCNA* normalized to *GFP* group. Each dot refers to a biological repeat. Data were presented as mean ± SEM. **$P < 0.01$ (*NTRK2-T1*+BDNF vs. *GFP*+BDNF, *NTRK2-T1*+BDNF vs. *NTRK2-FL*+BDNF), *** $P < 0.001$ (*NTRK2-FL*+BDNF vs. *GFP*+BDNF, *NTRK2-T1*+BDNF vs. *GFP*+BDNF, *NTRK2-T1*+BDNF vs. *NTRK2-FL*+BDNF), one-way ANOVA followed by Bonferroni's multiple comparisons (n = 3 repeats).

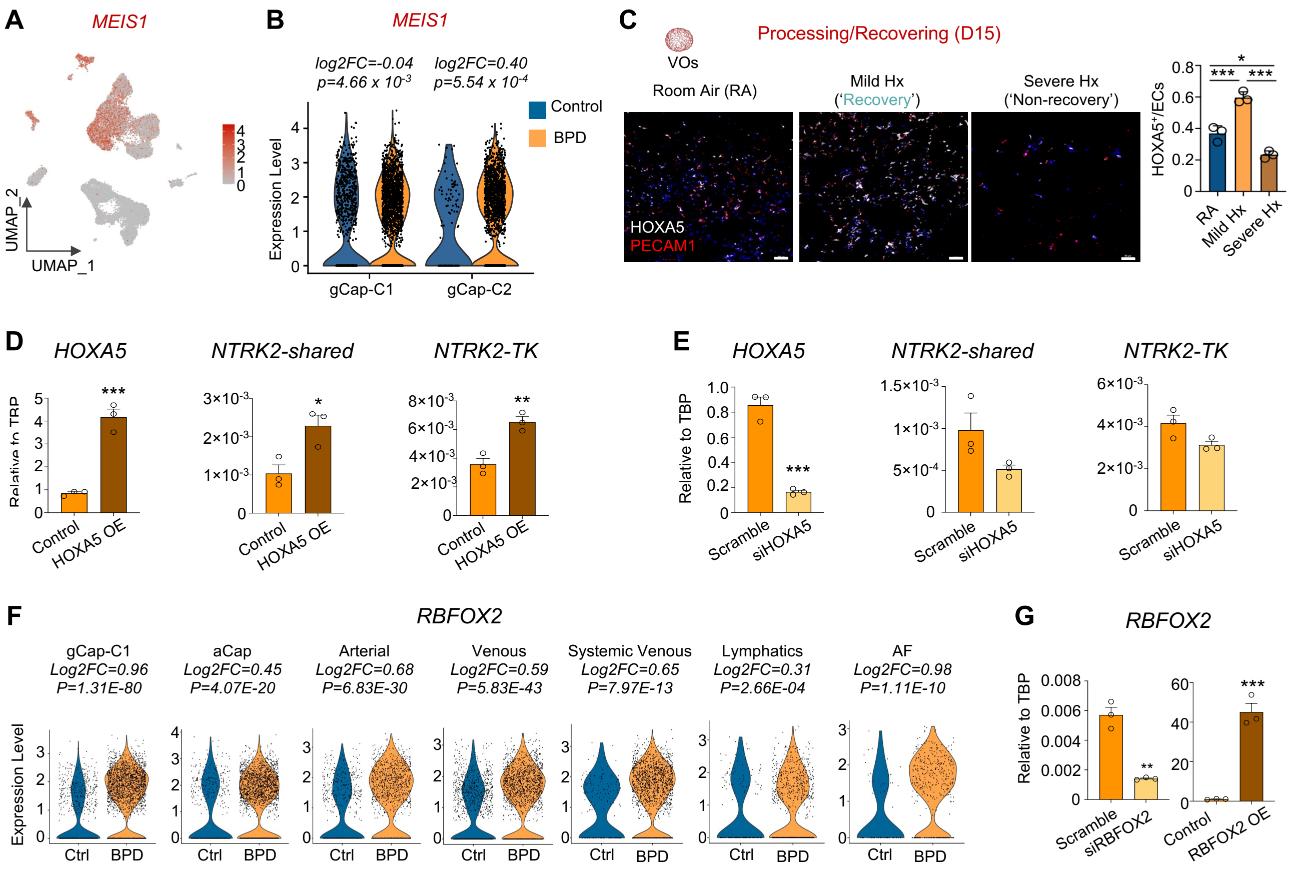


**Figure S7: RBFOX2 and HOXA5 mediated regulation of *NTRK2* isoforms.**

(**A**) UMAP visualization of expression of *MEIS1* in the 10x multiome profiling of BPD and control lung.

(**B**) Violin plot showing the expression of *MEIS1* in BPD vs. control of gCap-C1 and gCap-C2 cells in the multiome profiling. P values and fold change (FC) values were obtained from tests of gene expression in BPD vs. control cells within each cell type using Seurat FindMarkers function using two-tailed Wilcoxon rank sum test.

(**C**) Immunofluorescence staining of HOXA5 in Day15 VOs under different hyperoxia conditions according to figure 2J (*left*). HOXA5 (white), PECAM1 (red), DAPI (blue). Bar plot showing HOXA5$^+$ endothelial cells (ECs) among all ECs in different groups (*right*). Each dot represents one iPSC line. Data was represented as mean ± SEM. **P* < 0.05 (Severe Hx vs. RA), ****P* < 0.001 (Mild Hx vs. RA, Severe Hx vs. Mild Hx), one-way ANOVA followed by Bonferroni's multiple comparisons test (n = 3 iPSC lines). Scale bar, 50 µm.

(**D**) The expression of *HOXA5* and different isoforms of *NTRK2* after *HOXA5* overexpression. Each dot represents 1 biological repeat. Data were represented as mean ± SEM. **P* < 0.05, ***P* < 0.01, ****P* < 0.001 (vs. Control group), unpaired two-tailed *t* test (n = 3 repeats).

(**E**) The expression of *HOXA5* and different isoforms of *NTRK2* after *HOXA5* knockdown. Each dot represents one biological repeat. Data were represented as mean ± SEM. ****P* < 0.001 (vs. Scramble group), unpaired two-tailed *t* test (n = 3 repeats).

(**F**) Violin plot of *RBFOX2* expression in BPD vs. control cells of different cell types in the multiome profiling. Differential expression was performed comparing gene expression in control cells vs. BPD cells. P values and fold change (FC) values were obtained from tests of gene expression in BPD vs. control cells within each cell type using Seurat FindMarkers function using two-tailed Wilcoxon rank sum test.

(**G**) The expression of *RBFOX2* after knockdown or overexpression of RBFOX2 in human pulmonary microvascular endothelial cells (HPMVECs). Each dot represents one biological repeat. Data were represented as mean ± SEM. ** *P* < 0.01 vs. Scramble group, ****P* < 0.001 vs. Control group, unpaired two-tailed *t* test (n = 3 repeats).

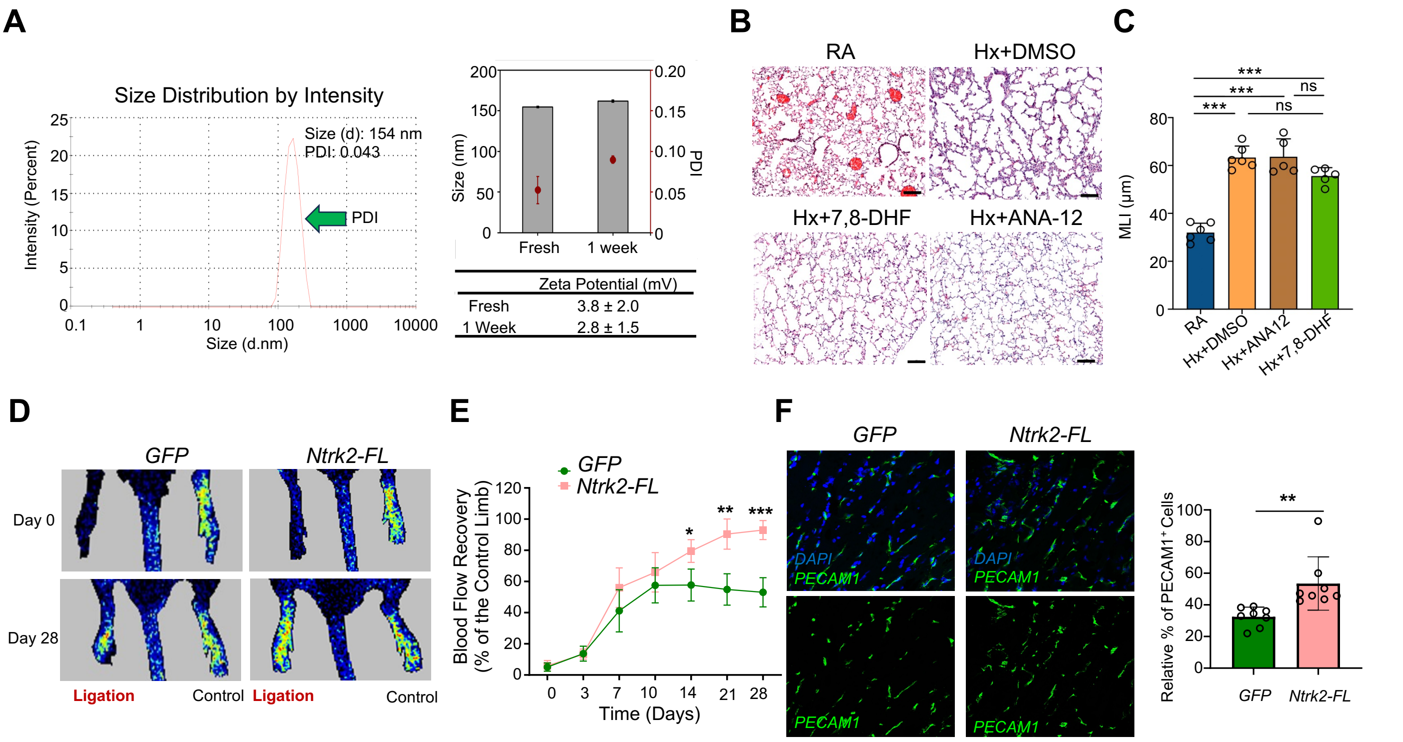


**Figure S8: Potential therapeutic applications of *Ntrk2* overexpression in pulmonary vascular regeneration and functional recovery in hindlimb ischemia.**

(**A**) Size distribution of synthesized LNP(left). Comparison of size, PDI and Zeta potential between fresh group (freshly made LNP) and 1 week group (stored for 1 week under -80 °C).

(**B**) Representative H&E staining image of P28 mouse lung under room air(RA), 90% hyperoxia treated with DMSO control (Hx+DMSO), 90% hyperoxia treated with 7,8-DHF (Hx+7,8-DHF) and 90% hyperoxia treated with ANA-12(Hx+ANA-12). scale bar, 100 μm. n= 6 mice.

(**C**) Quantification of MLI among groups in (B). Each dot represents one mice. ***$P < 0.001$ (Hx+DMSO group vs. RA group, Hx+ANA-12 group vs. RA group, Hx+7,8DHF group vs. RA group), one-way ANOVA followed by Bonferroni's multiple comparisons test (n = 6 mice).

**(D)** Representative images of blood flow to the ischemic and unoperated hindlimbs quantified by Laser Doppler system. Limb perfusion was monitored immediately after surgery, and 3, 7, 14, 21 and 28 days postoperatively.

**(E)** Line graph showing the blood flow in (D). Data were shown as mean ± SEM. n = 6 mice. *$P < 0.05$, **$P < 0.01$, ***$P < 0.001$ Ntrk2-FL vs. GFP, unpaired two-tailed *t* test (n = 6 mice).

**(F)** Fluorescence staining (left) and Bar plot (right) showing CD31 positive cells among all cells in mice with hindlimb ischemia surgery on day 28. Data were represented as mean ± SEM. **$P < 0.01$ (*NTRK2-FL* group vs. *GFP* group), unpaired two-tailed *t* test (n = 8 mice).
